# Perturbations in 3D genome organization can promote acquired drug resistance

**DOI:** 10.1101/2021.02.02.429315

**Authors:** Anna G Manjón, Daan Peric Hupkes, Ning Qing Liu, Anoek Friskes, Stacey Joosten, Hans Teunissen, Marleen Aarts, Stefan Prekovic, Wilbert Zwart, Elzo de Wit, Bas van Steensel, René H Medema

**Affiliations:** Oncode Institute, The Netherlands Cancer Institute, Plesmanlaan 121, 1066CX, Amsterdam, The Netherlands.; Division of Cell Biology, The Netherlands Cancer Institute, Plesmanlaan 121, 1066CX, Amsterdam, The Netherlands.; Division of Gene Regulation, The Netherlands Cancer Institute, Plesmanlaan 121, 1066CX, Amsterdam, The Netherlands.; Division of Oncogenomics, The Netherlands Cancer Institute, Plesmanlaan 121, 1066CX, Amsterdam, The Netherlands.

## Abstract

Acquired drug resistance is a major problem in the treatment of cancer. hTERT-immortalized, untransformed RPE-1 (RPE) cells can acquire resistance to taxol by derepressing the *ABCB1* gene, encoding for the multidrug transporter P-gP. Here we have investigated how the *ABCB1* gene is derepressed. We show that activation of the *ABCB1* gene is associated with reduced DNA methylation, reduced H3K9 trimethylation and increased H3K27 acetylation at the *ABCB1* promoter. In addition, we find that the *ABCB1* locus has moved away from the nuclear lamina in the taxol-resistant cells. This raises the question which of these alterations were causal to derepression. Directly modifying DNA methylation or H3K27 methylation had neither significant effect on *ABCB1* expression, nor did it promote drug resistance. In contrast, the disruption of Lamin B Receptor (LBR), a component of the nuclear lamina involved in genome organization, did promote the acquisition of a taxol-resistant phenotype in a subset of cells. Using CRISPRa-mediated gene activation, we could further substantiate a model in which disruption of lamina association renders the *ABCB1* gene permissive to derepression. Based on these data we propose a model in which nuclear lamina dissociation of a repressed gene allows for its activation, implying that deregulation of the 3D genome topology could play an important role in tumor evolution and the acquisition of drug resistance.

## Introduction

Chemotherapy, is one of the main pillars of cancer treatment. However, chemotherapeutic drugs loose efficacy over time due to acquired drug resistance^1,2^. This acquired drug resistance can be the result of genetic mutations, as exemplified by mutations in receptor tyrosine kinases that causes resistance to tyrosine kinase inhibitors^3,4^. Alternatively, drug resistance can arise through elevated gene expression of the drug target itself, or by altered expression of proteins involved in drug metabolism^5^. The cause of this altered gene expression can be a genetic mutation or amplification of one of its upstream regulators, but changes in gene expression can also be due to epigenetic changes^6,7^. Well known examples of these, are changes in DNA methylation that result in altered gene expression in cancer^8^. How exactly these changes are induced during the evolution of drug resistance is currently unclear.

Here we have investigated the process of gene activation in the evolution of drug resistance in non-transformed immortalized human cells in culture. We have used derepression of the *ABCB1* gene as our model system to study gene regulation and acquired drug resistance.

Extensive research has shown that the *ABCB1* gene (also known as multidrug resistance gene or MDR) encoding the P-glycoprotein (P-gP) drug-efflux pump, is upregulated in many cancers cells exposed to increasing doses of taxol and a variety of other chemotherapeutic drugs^9,10^. The contribution of P-gP to taxol resistance in patients is still debatable, with the possible exception of ovarian cancer, where it has been shown that taxol resistance correlates with increased ABCB1 expression^11^. In this same tumor type, *ABCB1* has been found fused with active promoters in taxol resistant samples^12,13^.

Prior studies have investigated the mechanisms of the *ABCB1* upregulation in cellular systems, and found that DNA-copy number amplifications of *ABCB1* locus can be linked to acquired chemoresistance^14^. Additionally, recent studies have shown that epigenetic alterations can also drive the upregulation of *ABCB1*. Particularly, several studies in taxol-resistant cancer cell lines demonstrated that loss of repressive marks of heterochromatin, such as DNA methylation, in the regulatory region was associated with active transcription of the *ABCB1* gene^15–18^.

Although prior reports suggest a role for the methylation status in *ABCB1* regulation, the influence of the higher-order chromatin structure on gene expression and drug resistance is not yet understood. In general, alterations in chromatin organization have been correlated to changes in gene expression^19–23^, and consequently, dysregulation of these may influence the functionality of the genome, leading to pathogenesis. It is well understood that the three-dimensional genome is maintained by a multilayer of structural units like chromosome territories, nuclear compartments, Topological Associating Domains (TADs) and Lamina Associated Domains (LADs). While chromosome compartments are proposed to be mediated by Condensin II and phase separation, TADs are often defined by CTCF and the cohesin complex^24–26^.

Several investigations have found alterations of the 3D genome involving TAD perturbations in cancer^26–28^ as well as in autoimmune diseases and limb malformations^29,30^. Furthermore, a recent study reported genomic CTCF-binding site mutations in 200 patient samples of colorectal cancer^31^. In addition to genomic organization in TADs, in the cell nuclei extensive chromatin regions are associated with the Nuclear Lamina (NL), which are mostly transcriptionally repressed^32,33^. This raises the question whether the NL could act as a repressive element for genes. Recent studies in *Drosophila* suggest that depletion of NL components alters gene expression of several chromatin regions, leading to defective cell differentiation^34–36^. However, in the context of drug resistance, it has not yet been examined whether 3D genome disorganization and detachment from the NL could be a potential mechanism of gene reactivation and consequently chemoresistance.

In order to explore novel mechanisms of gene re-activation and taxol resistance, we generated taxol-resistant cell lines derived from hTERT-immortalized, untransformed RPE-1 (RPE) cells. Consistent with our previous work^37^, we find that these cells become resistant to taxol through re-activation of the *ABCB1* gene. In taxol-sensitive cells, *ABCB1* is located in a LAD together with other inactive genes. We show that modifying chromatin marks by drug inhibition of DNA and histone-methyltransferase enzymes does not have a significant effect on the ABCB1 expression. In addition to the observed changes in chromatin modifications, we observe important changes of the 3D genome topology when comparing the taxol-sensitive versus the taxol-resistant lines, particularly in the NL interactions. Furthermore, the disruption of LBR, a NL component, is able to de-repress the locus leading to a taxol-resistant phenotype. Therefore, this research provides a new understanding, from a high-order chromatin perspective, of how cells may gain resistance to chemotherapeutics such as taxol.

## Results

### Transcriptional activation of *ABCB1* drives taxol resistance in RPE-TxR

In order to gain more insight in the processes that can lead to acquired drug resistance, we explored the molecular mechanism underlying *ABCB1* upregulation in the context of chemotherapy resistance. We made use of a previously described taxol-resistant cell line derived from hTERT-immortalized, untransformed RPE-1 cells obtained after prolonged exposure to increasing doses of taxol (RPE-Taxol Resistant, RPE-TxR)^37^. The generated cell line can proliferate under a taxol concentration 20-fold higher than the parental RPE-1 (RPE-Taxol Sensitive, RPE-TxS) (**Fig 1A and B**). Inhibition of the drug efflux pump P-gP by Tariquidar showed a re-sensitization of the RPE-TxR, indicating that Pg-P mediates resistance to taxol in this cell line (**Fig1A and B**)^37^. We independently generated new taxol-resistant RPE cell lines (TxR-3 and TxR-4) and confirmed that P-gP expression also conferred taxol resistance in these lines (**Sup Fig1A-B**). To interrogate whether enhanced P-gP protein expression was due to transcriptional activation of the *ABCB1* gene, we performed RT-qPCR analysis and observed that the mRNA level of *ABCB1* was increased in all of our clones (**Fig 1C, Sup Fig 1C**). In addition, single-molecule RNA FISH (smRNA FISH) revealed an increased number of active *ABCB1* Transcription Sites (TS) in RPE-TxR compared to RPE-TxS (**Fig 1D and E**). Taken together, we corroborate in three independently generated cell populations that the major mechanism underlying acquired taxol resistance in RPE-1 cells is through transcriptional activation of the *ABCB1* gene.

**Figure 1.**
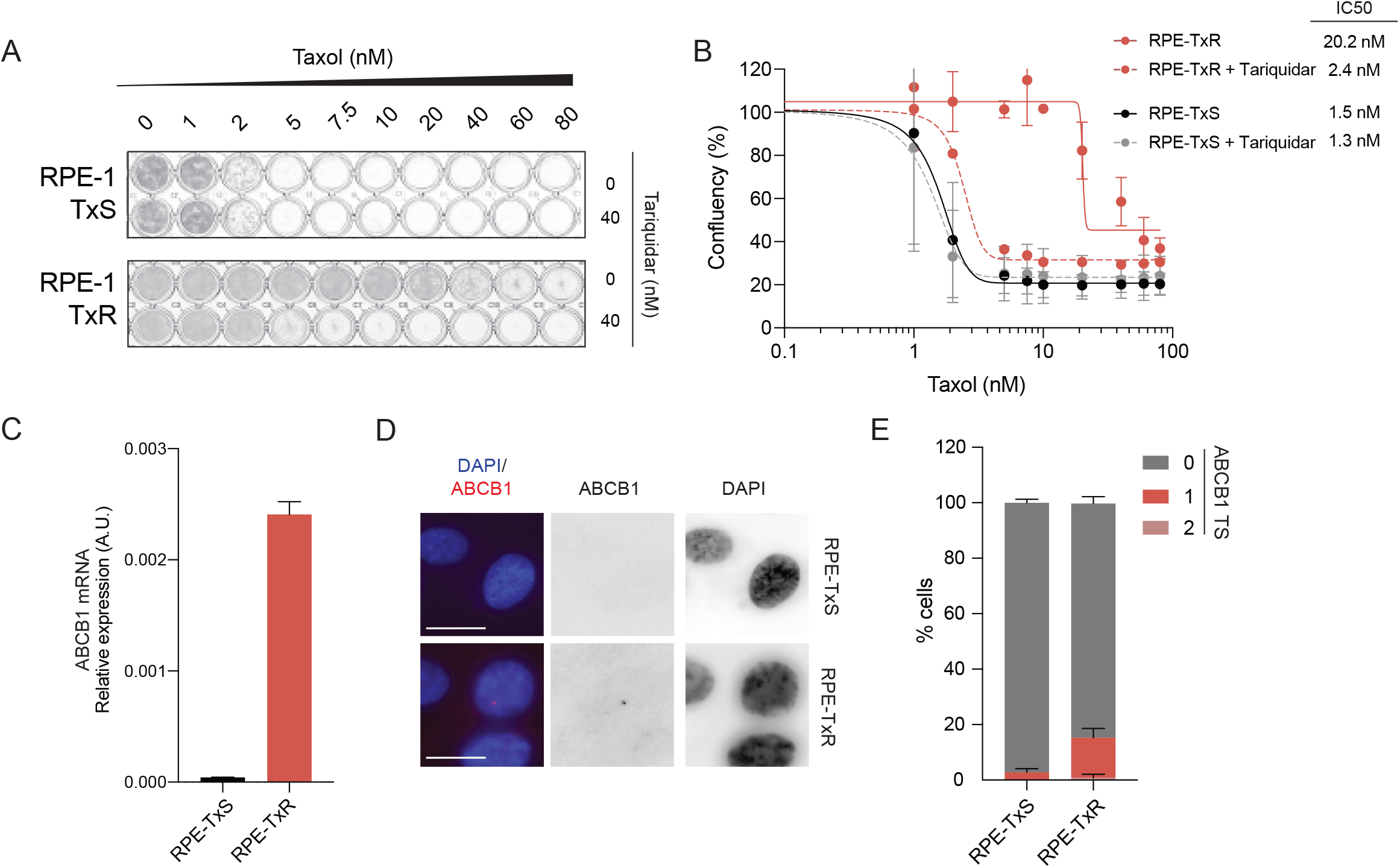
Transcriptional activation of *ABCB1* drives taxol resistance in RPE-TxR. **A)** Crystal violet staining of viability assay on taxol-naïve RPE-TxS and resistant RPE-TxR cell lines. **B)** Relative survival plots of the RPE-TxS and RPE-TxR cell lines. Error bars show the average +/− s.d. of two independent experiments and the calculated IC50. The curve was drawn from the log(inhibitor) vs response equation *Y=Bottom + (Top-Bottom)/(1+10^(X-LogIC50))*. **C)***ABCB1* mRNA levels determined by qRT-PCR and normalized to *GAPDH* expression levels, n=2. Error bars show the SD. **D)** Representative smRNA-FISH images of RPE-TxS and RPE-TxR for the *ABCB1* gene and DAPI. The images are projections of 0.5μm sections and a total 5μm in thickness. Scale bar, 15μm. **E)** Quantification of the number of *ABCB1* transcription sites (TS) found per cell, n=2, 60 cells per condition.

### *ABCB1* gene activation in RPE-TxR is associated with changes in chromatin modifications and DNA contacts at the ABCB1 locus

In order to understand the mechanism of upregulation of *ABCB1* in RPE-TxR cells, we first aimed to investigate whether *ABCB1* expression is accompanied by changes in chromatin modifications at the *ABCB1* locus. To this end, we analyzed histone marks and DNA methylation patterns by Chromatin and Methylated Immunoprecipitation (Ch-IP and MeDIP). We found that RPE-TxR lost repressive modifications (H3K9me3 and DNA methylation) and gained active marks (H3K27ac and H2AZ) in the promoter region of the *ABCB1* gene (**Fig 2A**), compared to RPE-TxS. Hi-C analysis demonstrated that *ABCB1* is found in a TAD together with two other genes, *ABCB4* and *RUNDC3B* (**Sup Fig 2A**). Interestingly, RNA-sequencing experiments showed, in addition to the 7-fold increase in the *ABCB1* mRNA levels, an upregulation of *ABCB4* and *RUNDC3B* in RPE-TxR (**Sup Fig 2B**). The same was seen in the additional independently generated RPE-1-derived taxol-resistant cell lines (TxR-3 and TxR-4) (**Sup Fig 2C-D**). Because changes in gene regulation are often associated with local changes in chromosome folding^38^, we performed Targeted Locus Amplification (TLA) in RPE-TxS and RPE-TxR. This strategy allows to selectively amplify and sequence DNA on the basis of the crosslinking of physically proximal sequences similarly to 4C-seq^39^. We identified changes in chromatin contacts of the *ABCB1* locus in RPE-TxR compared to RPE-TxS (**Fig 2B**). In RPE-TxS, *ABCB1* preferentially interacts with regions enriched for H3K9me3 and low for H3K36me3, associated with heterochromatin and transcriptionally active regions respectively^40,41^ (**Fig 2B**). However, in RPE-TxR, contacts also occurred in less enriched H3K9me3 domains. Moreover, new interactions with the promoters of the transcribed genes *SLC25A40*, *CROT, DMTF1* and *TMEM243* were observed, marked by H3K36me3 and H3K4me3 (**Fig 2B, Sup Fig 2E**). These new interactions were also enriched on H3K4me1, an enhancer-associated mark^42^, suggesting that the *ABCB1* gene could potentially be activated by proximal enhancers. Therefore, we conclude that chromatin marks undergo major changes at the *ABCB1* locus during the acquisition of taxol-resistance. This is also the case for *ABCB1* DNA interactions, suggesting that genes are more likely to interact with regions with similar chromatin nature.

**Figure 2.**
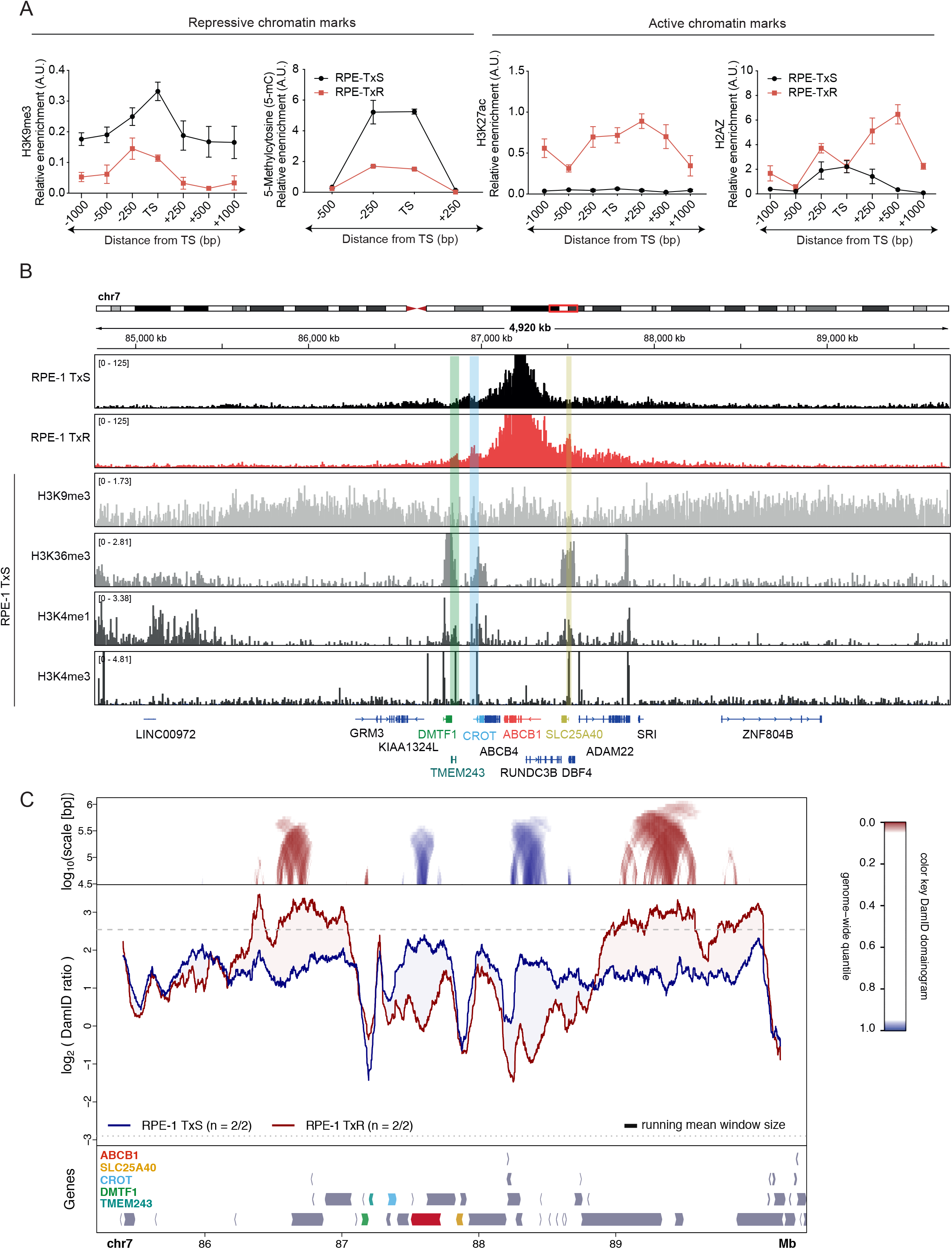
*ABCB1* gene activation in RPE-TxR is associated with changes in chromatin modifications and 3D genome. **A)** ChIP-qPCR of indicated chromatin and DNA methylation marks in the ABCB1 regulatory region for RPE-TxS and RPE-TxR. TS marks the transcription start site of the promoter. ChIP signal was normalized over input and a positive control specific for each mark, n=2. **B)** TLA analysis of the *ABCB1* gene in RPE-TxS (first row) and RPE-TxR (second row), (gene annotation hg19). Sequencing expanding 5000kb shows that regions immediately neighboring the *ABCB1* gene have higher coverage. 3^rd^ to 6^th^ rows: ChIP-sequencing tracks of indicated histone modifications in RPE-TxS cells expanding 5000kb from the *ABCB1* gene (gene annotation hg19). Color lines show new contacts formed in RPE-TxR with the indicated colored genes. **C)** Change is NL interactions of *ABCB1* and flanking regions in RPE-TxR compared to RPE-TxS. *Bottom panel:* gene annotation track (hg38) with indicated colored genes. *Middle panel:* DamID tracks of NL interactions in RPE-TxS (blue line) and RPE-TxR cells (red line). Data are the average of 2 independent replicates. Noise was suppressed by a running mean filter of indicated window size. Shading between the lines corresponds to the color of the sample with the highest value. Dotted lines mark the 5^th^ and 95^th^ percentiles of genome-wide DamID values. *Top panel*: domainograms; for every window of indicated size (vertical axis) and centered on a genomic position (horizontal axis), the pixel shade indicates the ranking of the change in DamID score (experimental minus control) in this window compared to the genome-wide changes in DamID scores across all possible windows of the same size. Blue: DamID score is highest in control samples; red: DamID score is highest in experimental samples.

### *ABCB1* gene activation in RPE-TxR is associated with detachment from the NL

As gene silencing has been linked to association with the Nuclear Lamina (NL)^43^, we also performed Lamin-DamID to study the *ABCB1*-NL interactions. We observed that in RPE-TxS, that the DamID signal intensity of the *ABCB1* locus is very high (**Fig 2C**, blue line), indicating that it is in a lamina-associated domain (LAD). In contrast, in RPE-TxR cells the DamID signal intensity is greatly reduced (**Fig 2C**, red line), indicating that a major NL detachment of the region containing *ABCB1* and its neighboring has taken place during the acquisition of drug resistance. Interestingly, *SLC25A40*, *CROT, DMTF1* and *TMEM243* are also found detached from the NL in RPE-TxS, suggesting that when *ABCB1* loses its interaction with the NL, it tends to interact with other inter-LAD (iLAD) genes, consistent with our TLA analysis (**Fig 2B**). In addition to this, we could also observe a possibly ‘compensatory’ movement of the regions further from the *ABCB1* locus, which increased NL contacts in the taxol-resistant cell lines (**Fig 2C**, red line). Interestingly, this phenomena has been previously reported in other loci^44^. Overall, these results indicate that a local rewiring of NL interactions occur in the *ABCB1* genomic region in the RPE-1 taxol resistant cells.

### Transition to taxol-resistance is not primarily driven by repressive chromatin modifications of *ABCB1* genomic locus

In order to test whether altering the chromatin modifications of the *ABCB1* locus is sufficient to de-repress *ABCB1* in RPE-TxS, we made use of different drugs to perturb the epigenetic landscape. The addition of 5-aza deoxycytidine (5-AZA) for 24h was able to reduce the levels of DNMT1, the enzyme responsible for DNA methylation deposition (**Fig 3A**). A similar trend for the levels of H3k27-trimethylation occurred when treating cells with GSK126 (EZH2 inhibitor), that interferes with H3K27me3 deposition (**Fig 3A**). Under these treatments, we performed RT-qPCR in RPE-TxS to check *ABCB1* expression levels. We observed that both drugs were unable to induce transcription of the *ABCB1* gene (**Fig 3B**). Thus, altering the levels of the H3K27-methyltransferase EZH2 or the DNA methyltransferase DNMT1 is not sufficient to derepress the *ABCB1* gene.

**Figure 3.**
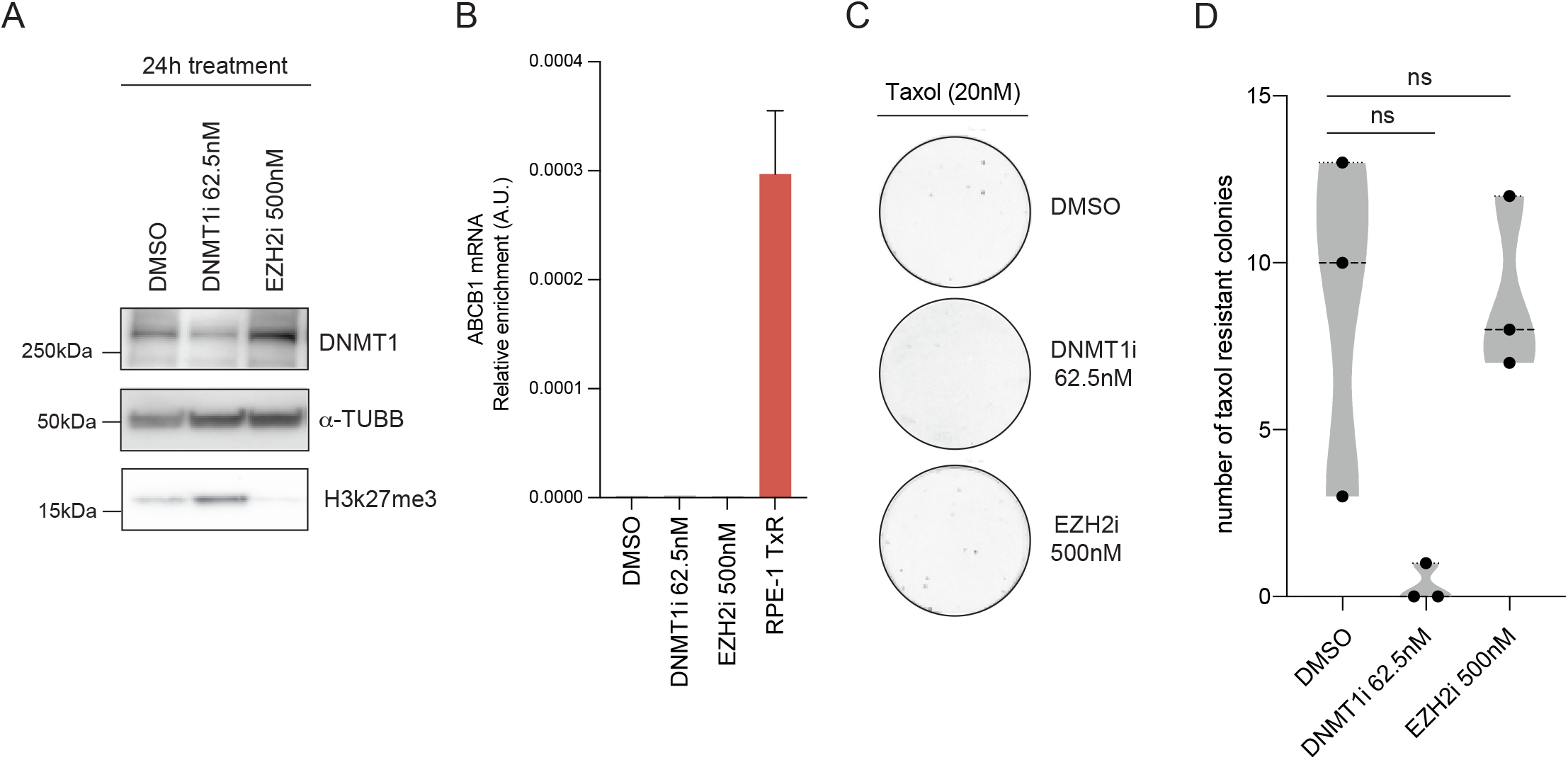
Transition to taxol-resistance is not primarily driven by repressive chromatin modifications of *ABCB1* genomic locus. **A)** Western Blot showing the levels of the chromatin proteins and controls (α-TUBB) upon treatment with the indicated epigenetic drugs with for 24h. **B)***ABCB1* mRNA levels determined by qRT-PCR and normalized to *GAPDH* expression levels upon drug addition and RPE-TxR as a control for *ABCB1* expression, n=2. Error bars show the SD. **C)** Crystal violet staining of colony formation assay under 20nM of taxol and the corresponding chromatin drug in RPE-1 iCut WT cells. **D)** Quantification of the number of taxol resistant colonies from B. Black dots show an independent biological replicate. ns, p>0,05, Mann-Whitney test.

We next asked if altering H3K27-trimethylation or DNA methylation at the *ABCB1* promotor is sufficient to precondition the locus for derepression. To this end, we performed colony formation assays using a combination of the epigenetic drugs and taxol. For the chromatin drugs we determined a dose that did not induce a proliferation defect (**Sup Fig 3A-B**). We pre-treated RPE-TxS cells with DNMT1i or EZH2i for 24h followed by an over-night co-treatment with 20nM taxol. Next morning the epigenetic drugs were washed out and only 20nM taxol was present for 15 days. Neither the DNMT1 nor EZH2 inhibitor were able to increase the number of taxol-resistant colonies (**Fig3C-D**). In fact, DNMT1i in combination with taxol led to a decrease in taxol-resistant colonies compared to the DMSO control (**Fig3C-D**). To boost the drug efficacy, we treated RPE-TxS cells for 72h with a higher dose of DNMT1i and maintained the same EZH2i dose. In addition, we included the H3k9me2 methyltransferase G9a inhibitor BIX01294. We observed a protein decrease on DNMT1, H3K27me3 and H3k9me2 when treating cells with DNMT1i, EZH2i and G9ai respectively or in combination (**Sup Fig 3C**). Moreover, an overall increase of the active mark H3k27ac and a decrease of the repressive mark H3k9me2 was detected by immunofluorescence in the cell nucleus (**Sup Fig 3D-E**). However, *ABCB1* mRNA levels quantified by qPCR remained similar to the DMSO-treated condition (**Sup Fig 3F**). Therefore, these data suggest that the disruption of chromatin-modifying enzymes by drug inhibition is unable to trigger activation of *ABCB1* gene transcription in RPE-1 cells, and thereby remain taxol-sensitive.

We next investigated whether potential upregulation of transcription factors (TFs) in RPE-TxR cells could be responsible for initiation of *ABCB1* gene expression, and thereby change local chromatin modifications and 3D genome organization at the *ABCB1* locus. We first performed motif scan to identify the potential TFs binding to the promoters of the two *ABCB1* isoforms. Subsequently, we hypothesized that gain of taxol resistance may be caused by aberrant expression of some of these TF interactors, and therefore we identified all the differentially expressed TF binders of the two promoters in RPE-TxR compared to RPE-TxS using mRNA sequencing (**Sup Fig 4A**). To further narrow down our searching, we speculated that the TFs responsible for the *ABCB1* derepression may potentially play an activation role for other upregulated genes in the resistant cells. Hence, we also performed a motif analysis for the promoters of all the upregulated genes in RPE-TxR in order to identify general promoter activators in the resistant cell line. We mainly found significantly enriched motifs belonging to the POU and LHX TF homeodomain family (**Sup Fig 4B-C**). This implies that these TFs may potentially be involved in the upregulation of many genes on the RPE-TxR cell lines, including *ABCB1*. To test this, we overexpressed *POU3F2*, *LHX6* and *ZIC5,* which showed a clear upregulation in the resistant cells (**Sup Fig 4A**), in the taxol sensitive parental RPE cells. To that aim, we used the Cas9-VP64-transcription activation system (CRISPRa) to assess whether this would recapitulate *ABCB1* activation in the resistant cell line. Even though we observed by RT-qPCR a significant increase of mRNA expression of the three TFs, similar to the level of upregulation in RPE-TxR (**Sup Fig 4D**), this did not result in a taxol-resistant phenotype (**Sup Fig 4E-F**). More importantly, downregulation of these TFs in RPE-TxR did not perturb the taxol-resistant phenotype (**Sup Fig4 G-I**), clearly indicating that *POU3F2*, *LHX6* and *ZIC5* are not required for expression of the *ABCB1* gene in RPE-TxR cells.

**Figure 4.**
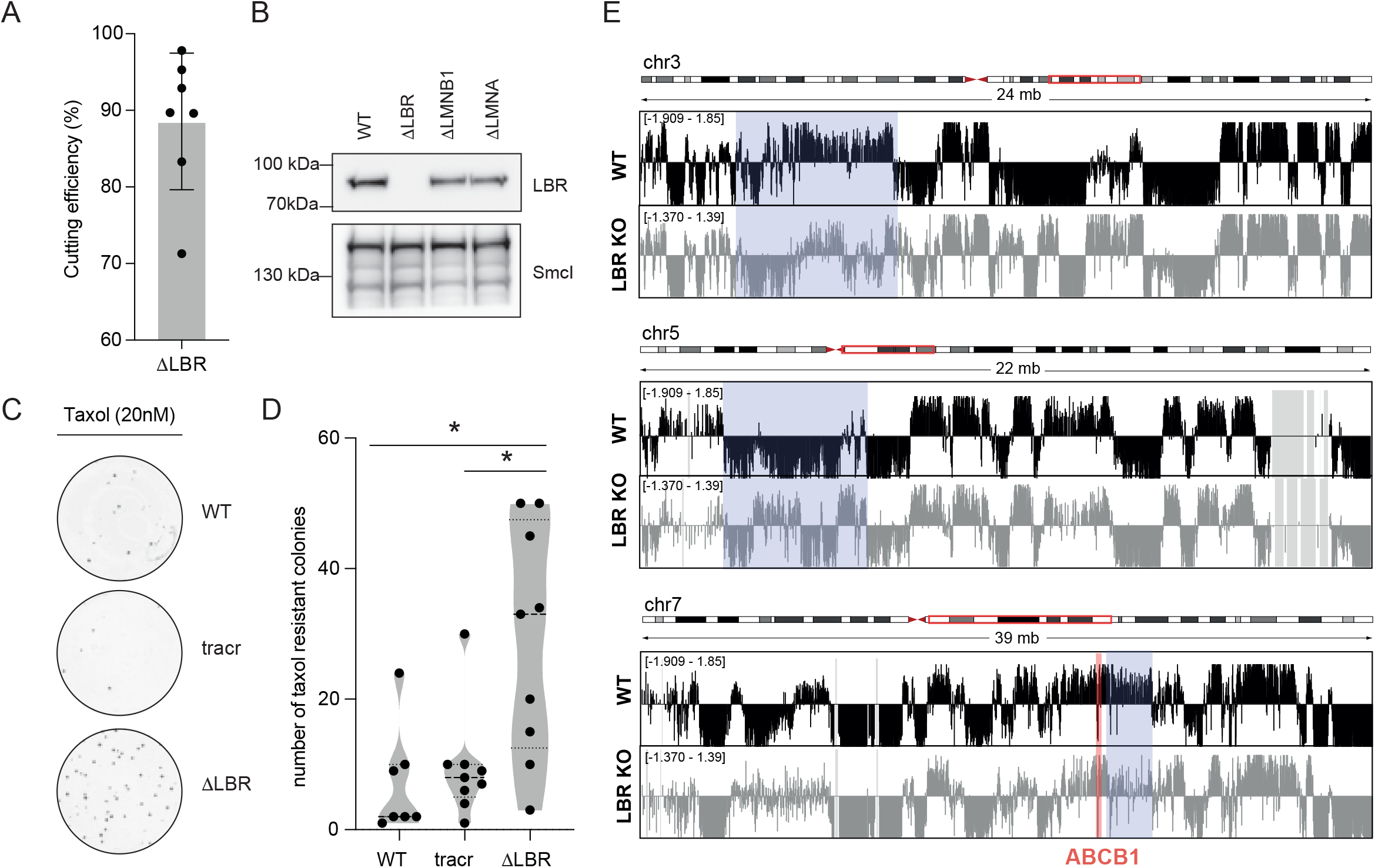
LBR depletion facilitates acquisition of taxol resistance. **A)** Percentage of disrupted sequence (cutting efficiency) in RPE-1 iCut cells transfected with crRNAs targeting the *LBR* gene and using TIDE analysis. Black dots show an independent biological replicate. Error bars show the SD. **B)** Western Blot showing the levels of *LBR* and control (SMC1) proteins 7 days after transfection of *LBR, LMNA or LMNB1* crRNAs. **C)** Crystal violet staining of colony formation assay under 20nM of taxol in RPE-1 iCut WT cells, transfected with only tracrRNA or tracrRNA and crRNA-*LBR*. **D)** Quantification of the number of taxol resistant colonies under 20nM of taxol in crRNA-*LBR* compared to WT and tracr only. Black dots show an independent biological replicate, *p<0,05, Mann-Whitney test. **E)** Change is NL interactions in *LBR* KO compared to RPE-TxS (WT) analyzed by DamID. Positive values indicate NL interactions, negative values indicate NL detachment. Three regions of the genome are shown. Blue: differences observed between WT and *LBR* KO. Red: *ABCB1* genomic region.

### *ABCB1* upregulation in RPE-TxR is not caused by direct activation of the promoter by trans-acting factors

We next wondered whether RPE-TxR cells upregulated additional TF that could lead to the activation of the *ABCB1* promoter. Therefore, to further exclude TF activation as the initial trigger for *ABCB1* gene activation, we carried out a luciferase reporter assay to assess the *ABCB1* promoter activity in RPE-1 taxol-sensitive (TxS) and taxol-resistant (TxR3-4) cells. To this end, the *ABCB1* promoter was cloned in a pGL3-basic vector followed by transfection into RPE-TxS or RPE-TxR. If similar luciferase activity was observed between cell lines, that would indicate that there are not differentially expressed trans-acting factors that lead to *ABCB1* promoter activation. However, if there is an increase of luciferase activity in RPE-TxR, a trans-acting factor may be upregulated therefore inducing *ABCB1* promoter activation. Activity of the ABCB1 promoter was relatively low compared to the pGL3-promoter plasmid, but more importantly, we did not observe an increase of luciferase activity in taxol-resistant cells compared to taxol-sensitive (**Sup Fig 5A**). This suggests that RPE-1 TxR cells do not have a distinct transcriptional program or differentially expressed TFs which could activate the *ABCB1* promoter. Instead, the 3D genome topology may be the determining factor for the *ABCB1* expression. Therefore, we hypothesized that NL detachment observed in RPE-TxR potentially could be a first step towards acquired drug resistance, subsequently allowing recruitment of available TFs leading to transcription activation of the *ABCB1* gene.

To further support the impact of NL in the regulation of *ABCB1*, we measured the *ABCB1* transcription levels in its native chromatin environment and outside of this context. We obtained these data from myelogenous leukemia K562 cells^32^. We used *GRO-cap* (global run-on sequencing with 5′cap selection) data as a measure of nascent RNA in native chromatin context.

In order to detect transcription outside the chromatin context we used the plasmid-based assay *SuRE* (Survey of Regulatory Elements)^45^. *ABCB1* exhibited a low *GRO*-cap activity and higher SuRE signal, suggesting that it is repressed by its native chromatin environment but can be activated when transcription activators have access to the regulatory region (**Sup Fig 5B**). All together, these results suggest that in RPE-1 WT cells, *ABCB1* is located in a repressive chromatin environment but has the ability to activate transcription if removed from this context.

### LBR depletion facilitates acquisition of taxol resistance

To further understand the importance of NL components in *ABCB1* gene expression we generated different knock-outs (KOs) of NL proteins using CRISPR-Cas9 technology in RPE-1 Cas9 cells (RPE-1 iCut)^46^. We obtained a high cutting efficiency of the Lamin B Receptor (*LBR*) gene in a polyclonal cell population (**Fig 4A**). Moreover, we could confirm by western blotting that *LBR* was depleted effectively (**Fig 4B**). 7 days after the KO generation we performed colony formation assays using 20nM of taxol. Upon *LBR* depletion, we observed an increase in the number of taxol-resistant colonies in multiple independent experiments (**Fig 4C-D**). Interestingly, the *ABCB1* mRNA levels were not increased in the polyclonal population (**Sup Fig 6A**). This implies that the loss of *LBR* can facilitate derepression of the *ABCB1* gene when cells are exposed to taxol. Clearly, loss of *LBR* alone is not sufficient for full derepression of the *ABCB1* gene, because we find that only a fraction of cells in the population acquires taxol-resistance. Moreover, the absolute number of taxol-resistant colonies we obtained varied from experiment to experiment, suggesting the importance of other factors in activating the *ABCB1* gene. Nevertheless, the number of taxol-resistant clones is significantly higher in *LBR* knock-out cells than what we observe in the parental lines. Complete depletion of Lamin B1 (*LMNB1*) or Lamin A/C (*LMNA*), structural and supporting components of the NL, had no obvious effect on taxol resistance (**Sup Fig 6B-E**).

In order to investigate whether depletion of *LBR* induced *ABCB1* upregulation in other *in vitro* models, we performed RNA interference experiments in various cancer cell lines. We selected a Triple Negative breast cancer (TNBC) cell line (MDA-MB-231), a head-and-neck squamous cell carcinoma (FaDu) and a lung adenocarcinoma cell line (A549). Using RT-qPCR analysis we found that MDA-MB-231 and FaDu had slightly higher *ABCB1* mRNA levels than RPE-1 cells. In contrast, the *ABCB1* mRNA levels detected in A549 were considerably increased (**Sup Fig 6F**). Depletion of *LBR* by siRNA led to a decrease of *LBR* protein levels 48h post-transfection in all cell lines (**Sup Fig 6G**). After 48h, colony formation assays under different concentrations of taxol for each of the cell lines were performed. As control, we confirmed that depletion of *LBR* by siRNA led to an increase in number of taxol-resistant colonies in RPE-1 cells (**Sup Fig6 H-I**). As expected, based on the high level of *ABCB1* expression, A549 cells were resistant to high levels of taxol, and depletion of *LBR* had minimal effects (Sup Fig.5H-I). The effect of *LBR* depletion in MDA-MB-231 also resulted in increased numbers of taxol-resistant colonies, similar to what we observe in RPE-1 cells (**Sup Fig 6H-I**). *LBR* depletion in FaDu cells resulted in a decrease of taxol-resistant colonies (**Sup Fig 6H-I**). These data imply that loss of *LBR* can prime *ABCB1* for derepression in some cells, but additional factors are required to achieve derepression.

To explore the reorganization of the LAD landscape that takes place upon *LBR* depletion we performed Lamin-DamID in the polyclonal RPE-1 *LBR* KO cells. Overall, the LAD landscape of the parental RPE-1 cells was largely retained in the *LBR* KO cells, and only a subset of LADs was clearly altered. Detachment of the *ABCB1* locus from the NL was not detected in this polyclonal population, but a decrease on NL interactions was seen the neighboring regions (**Fig 4E,** bottom panel). This change could destabilize the NL interactions of the locus and render the *ABCB1* locus more permissive for derepression. Alternatively, the effect of *LBR* depletion on NL interactions is not uniform across the entire population, causing the *ABCB1* locus to detach from the NL in a small subset of cells only. Finally, it is also possible that loss of *LBR* does not change the contacts of ABCB1 with the NL, but instead causes a reduced repressive potential of the NL. In support of the latter model, *LBR* was previously found to interact with the repressive protein HP1.^47,48^

Based on these data we propose that in RPE-1 cells, and possibly across various other *in vitro* models, *LBR* may act as a regulator of the *ABCB1* gene expression and its depletion can contribute to acquired taxol resistance. Additionally, these data suggest that NL-association may act as a critical threshold that needs to be overcome in order to derepress a gene, and as such loss of lamina-association might be a first step in the process of transcriptional derepression.

### Transcription-driven CRISPRa activation of neighboring genes can detach *ABCB1* from the NL and lead to taxol resistance

To further explore the role of NL in ABCB1 regulation, we examined whether NL detachment would lead to ABCB1 gene activation. It has been previously described that the CRISPRa induces detachment of genes from the NL, and in some instances this also causes detachment of flanking genes^44^. We therefore attempted to detach ABCB1 from the NL by activation of its neighboring genes. We used CRISPRa to specifically activate the promoter of *ABCB1*, *ABCB4* or *RUNDC3B* or a combination of the latter two (**Fig 5A**). Next, we performed Lamin-DamID to map NL interactions (**Fig 5B-C, Sup Fig 7A-B**). We observed that in control cells, *ABCB1* is located at the NL, together with the *ABCB4* and *RUNDC3B* genes (**Fig 5A-B, Sup Fig 7A-B**, blue lines). As expected and showed in previous research^44^, upon CRISPRa single gene activation local NL detachment was detected in the regulatory regions and most of the transcription units of these genes (**Sup Fig 7A-B**, red line). Strikingly, simultaneous activation of *ABCB4* and *RUNDC3B* caused not only detachment of these two genes, but also of *ABCB1* (**Fig 5C**, red line). Next, we asked whether this was accompanied by upregulation of *ABCB1* expression. We observed that transcription activation of *ABCB1* by CRISPRa led to an expected increase of mRNA of *ABCB1* (**Fig 5D**). Surprisingly, activating *ABCB4*, *RUNDC3B* or the combination via CRISPRa also triggered the activation of *ABCB1* (**Fig 5D, Sup Fig 7C**), and was accompanied by an increase in occurrence of taxol-resistant colonies (**Fig 5E, Sup Fig 7D-G**). We next performed ChIP-qPCRs on the *ABCB1* regulatory region and observed a decrease in the H3K9me3 signal in both CRISPRa-*ABCB1* and CRISPRa *ABCB4*-*RUND3CB* compared to the CRISPRa parental cell line (**Fig 5F**, left). However, even though CRISPRa-*ABCB1* presented an enrichment of H3k27ac in the *ABCB1* promoter, the combination of *ABCB4* and *RUNDC3B* did not show this (**Fig 5F**, right). To rule out the possibility that the *ABCB1* transcription initiation by *ABCB4* and *RUNDC3B* was a consequence of cross-activation of the *ABCB1* promoter, instead of a NL-detachment effect, we generated new sgRNAs targeting upstream and downstream of the *ABCB1* regulatory regions (**Sup Fig 7H**). We could confirm that these sgRNAs, even though in the same TAD as *ABCB1*, could not initiate transcriptional activation, as shown by RT-qPCR (**Sup Fig 7I**). Therefore, we could conclude that *ABCB1* transcription is linked to loss of H3K9me3 and NL detachment potentially caused by activation of *ABCB4* and *RUNDC3B* and not due to cross-activation of the sgRNAs.

**Figure 5.**
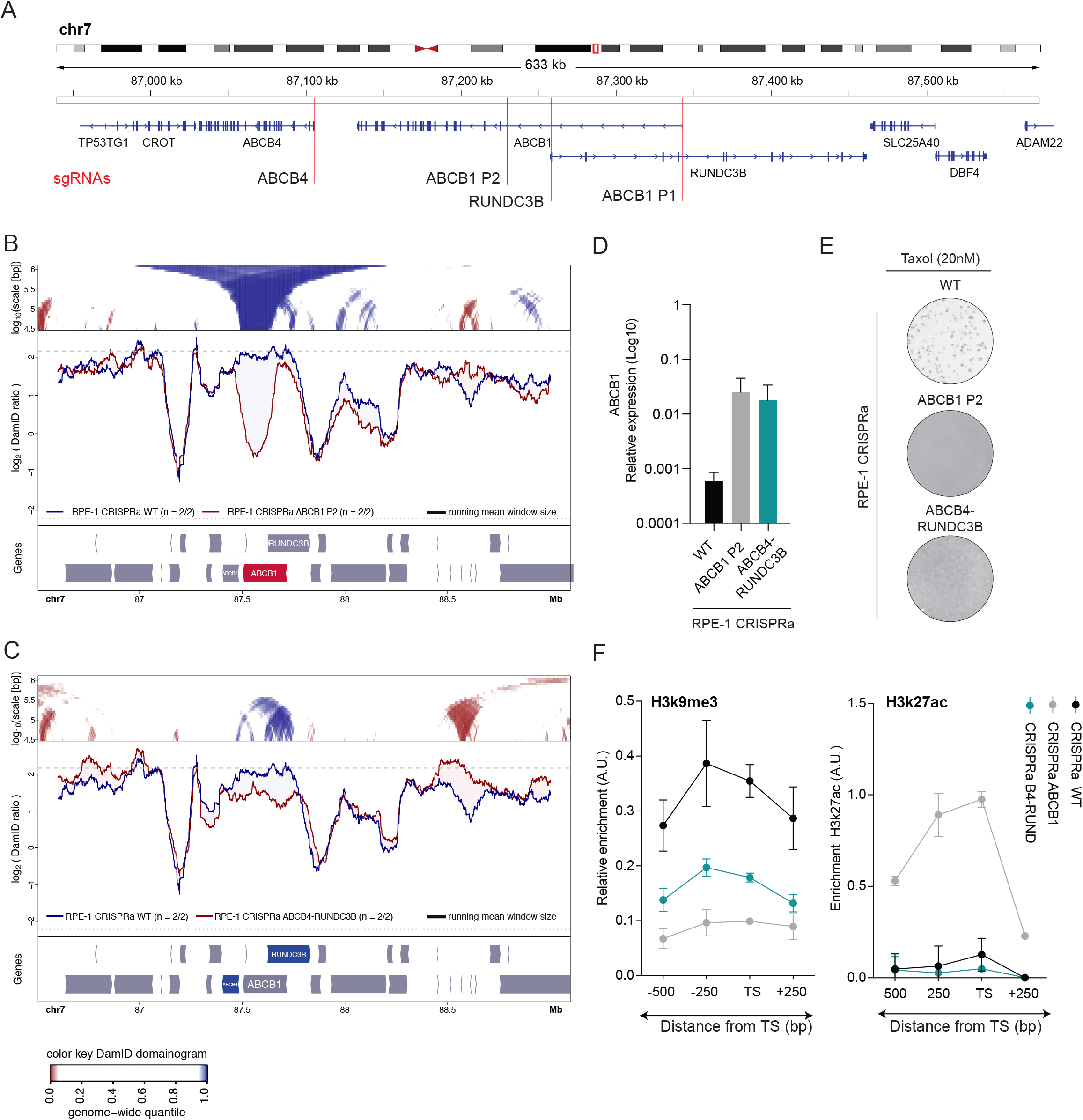
Transcription-driven CRISPRa activation of neighboring genes can detach *ABCB1* from the NL and lead to taxol resistance. **A)** Schematic representation of the Chr7q21.12 region indicating the locations where the sgRNAs were targeting for CRISPRa *ABCB1*, *ABCB4* or *RUNDC3B* activation. Two regions were independently targeted to upregulate ABCB1: P1 (proximal promoter, 6 sgRNA were used) and P2 (internal promoter, a single sgRNA was used). **B)** Local NL detachment caused by *ABCB1* gene activation by CRISPRa in RPE-1 cells. **C)** Local NL detachment caused by simultaneously *ABCB4* and *RUNDC3B* gene activation by CRISPRa in RPE-1 cells. **D)***ABCB1* mRNA levels determined by qRT-PCR and normalized to *GAPDH* upon CRISPRa activation of *ABCB1* (P2) or combination of *ABCB4* and *RUNDC3B*, n=2. Error bars show the SD. **E)** Crystal violet staining of viability assay on CRISPRa cell lines upon activation of *ABCB1* (P2) and the combination of *ABCB4* and *RUNDC3B*. **F)** ChIP-qPCR of H3K9me3 (left) and H3K27ac (right) in the *ABCB1* regulatory region for CRISPRa WT, *ABCB1* or the combination of *ABCB4* and *RUNDC3B* (B4-RUND). TS marks the Transcription Start Site of the promoter. ChIP signal was normalized over input and a positive control specific for each mark, n=2.

## Discussion

In this study we describe a novel mechanism by which cells can upregulate *ABCB1*, a gene involved in taxol resistance. Our data provide the first direct link between 3D genome reorganization and drug resistance. We have shown that taxol resistance of RPE-TxR cells can be entirely attributed to the activity of the P-gP drug efflux pump^37^. In RPE-TxR, *ABCB1*, the gene encoding for P-gP, is upregulated through transcriptional activation. This transcriptional activation coincides with an enrichment of active histone marks and a depletion of repressive marks in the chromatin environment of the *ABCB1* promotor. However, directly altering the chromatin landscape in RPE-TxS cells by drug inhibition of chromatin regulators did not lead to initiation of *ABCB1* expression. In addition to the altered chromatin modifications in the promotor region, we noted a clear detachment of the *ABCB1* locus from the NL in the taxol-resistant cells. In conjunction with that, disruption of the *LBR*, a key NL protein, led to enhanced acquisition of drug-resistance, implying that NL detachment can prime the *ABCB1* locus for gene activation.

### Role of histone modifications and DNA methylation in the ABCB1 locus

*ABCB1* gene regulation is thought to be driven by DNA methylation^49^. Some studies have shown that low DNA methylation status of the *ABCB1* promoter is linked to gene activation^15,16^. However, other studies were unable to confirm these findings^17,18^. Here we show that there is a switch from inactive to active chromatin in the *ABCB1* promoter in RPE-TxR cells, as well as a change in DNA methylation pattern. Depletion of the DNA methyltransferase DNMT1 in RPE-TxS cells did not directly alter *ABCB1* gene expression or taxol sensitivity. The same was observed when inhibiting the H3K27 methyltransferase EZH2, suggesting that the active chromatin environment observed in the *ABCB1* promoter region in RPE-TxR cells may be secondary to gene activation during the process of transcriptional derepression.

### How depletion of LBR may de-repress ABCB1

Studies in *Drosophila* have found that depletion of lamins can lead to de-repression of NL-associated genes^35,50^. Here, we found that *ABCB1* is partially activated upon depletion of LBR but not lamins. We speculate that depletion of *LBR* may lead to leaky *ABCB1* gene expression in at least two different ways. In one model, loss of *LBR* may cause stochastic detachment of *ABCB1* from the NL. In mouse and human cells LBR has been implicated in anchoring heterochromatin to the NL^51–53^. In our study, the frequency of heterochromatin detachment after LBR depletion may be too low to be detectable by DamID. However, if stable contact with the NL is essential for robust repression of *ABCB1*, then occasional detachment could account for the stochastic occurrence of taxol-resistant clones in *LBR*-depleted cells. Indeed, NL interactions can be intrinsically stochastic, and the NL contact frequency is inversely linked to gene expression^54,55^. This may explain why only a small proportion of cells acquires taxol resistance. In a second model, depletion of *LBR* may not affect the *ABCB1* – NL contact frequency, but rather may compromise the repressive potential of the NL. *LBR* may play a direct role in this repression, e.g. through its interaction with HP1^47^, or indirectly by controlling the protein composition of the NL. This partially defective repression in *LBR*-depleted cells could then allow for emergence of taxol-resistant clones. In both models, interactions of *ABCB1* with the NL contribute to its repression.

### Forced detachment of ABCB1 from the NL coincides with gene activation

The generation of CRISPRa cell lines targeting *ABCB4* and *RUNDC3B* allowed for detachment of *ABCB1* from the NL, and we find that this is associated with *ABCB1* gene activation. This further suggests a causal effect between NL detachment and *ABCB1* gene activation. However, we cannot fully rule out that the activated *ABCB4* and *RUNDC3B* promoters act as enhancers of the nearby *ABCB1* promoter, because enhancer activity has been observed for many promoters^56^. Interestingly, a decrease of gene expression has previously been observed by tethering chromosomes to the nuclear periphery^57–59^. Interestingly, recent research has shown that intrinsic features of promoters influence their sensitivity to the repressive LAD environment^32^. According to this study, *ABCB1* promoter is classified as repressed in K562 cells and thereby to have the potential to be activated if taken out from their native repressive LAD environment.

### Celltype-specific roles of LBR and lamins

We find that depletion of *LBR*, but not Lamin A/C, or B, can render the *ABCB1* locus permissive to gene activation. In another study, Lamin A/C together with *LBR* were shown to be involved in tethering heterochromatin to the nuclear periphery during development^53^. Interestingly, a recent study shows that loss of Lamin B1 leads to detachment of LADs together with global chromatin re-distribution and de-compaction, supporting the idea that NL have a role in chromatin dynamics and potentially in gene regulation^61^. Our results show that only *LBR* depletion has a positive effect on the induction of taxol resistance in RPE-1 cells. This could be because in differentiated cells, NL components may have different relevance on gene repression. Certainly, LBR may have celltype-specific effects, as we observed with the depletion of *LBR* in the various cancer cell lines.

Taken together, we propose that acquisition of taxol resistance in RPE-1 cells requires detachment of the *ABCB1* locus from the nuclear lamina as a priming event. When this priming event is followed by gene activation, this could induce changes to the local chromatin state that might help to keep the locus detached from the NL. Whether lamina detachment is the most critical step in the derepression of an inactive gene likely depends on the contribution of lamina-association in the regulation of gene expression of a given gene. One could envision that 3D genome rearrangements are an important priming step in the activation of a gene that is tightly associated to the NL, while activation of a TF is more likely to be the crucial event for activation of genes that display are more relaxed lamina-association.

## Acknowledgments

We thank the NKI Genomics for excellent support, and members from our laboratories for inspiring and helpful discussions. This work was supported by NWO Zwaartekracht (58588) (to R.H.M.) and by NIH Common Fund “4D Nucleome” Program grant U54DK107965 (to B.v.S.). The Oncode Institute is partly supported by KWF Dutch Cancer Society.

## Author contributions

Conceived and designed study: A.G.M., R.H.M.

Experiments: A.G.M., D.P.H, N.Q.L., A.F., S.J., H.T., M.A.

Data processing, bioinformatics: N.Q.L., S.J., E.d.W., B.v.S.

Discussion and interpretation of data: A.G.M., S.P., R.H.M., B.v.S.

Manuscript writing: A.G.M., R.H.M., B.v.S., with input from all authors.

## Materials and Methods

### Cell lines and cell culture conditions

hTert-immortalized retinal pigment epithelium (RPE-1) and derived cell lines were maintained in DMEM/F-12 + Glutamax (Gibco, Life Technology) supplemented with 1% penicillin/streptomycin and 6% fetal bovine serum (FBS, S-FBS-EU-015, Serana). A549 cancer cell lines were grown in Advanced RPMI 1640 (Gibco, Life Technology) supplemented with 1% penicillin/streptomycin, 1% sodium pyruvate, 2% HEPES buffer and 10% fetal bovine serum. MDA-MB-231 and FADU cell lines were maintained in DMEM (Gibco, Life Technology) supplemented with 1% penicillin/streptomycin, 1% sodium pyruvate, 2% HEPES buffer and 10% fetal bovine serum. All cell lines were routinely checked for mycoplasma.

### Drug treatments

Drugs were dissolved in DMSO and prepared at stock concentrations before usage at varying final concentrations as indicated in each figure. For the 24h assay, cells were treated for 24h with the specific epigenetic drug dose, adding 20nM of taxol overnight followed by a wash out of the drugs and subsequently addition of 20nM taxol again for 15 days. For the epigenetic drug treatment combination (Combo), 250nM of 5-Aza-2’-deoxycytidine, 150nM of GSK126 and 2μM of BIX-01294 were used.

### Luciferase Assay

The *ABCB1* promoter was cloned in a pGL3-basic (Promega) vector (pGL3-Basic Vector GenBank® Accession Number U47295). The *ABCB1* internal promoter region (1kb) was PCR amplified from RPE-1 genomic DNA and inserted downstream of the luciferase reporter gene. The primers used were: gatcAAGCTTCATTAGCCAAATGCATGAGC (FWD) and GATCGGTACCTGGAAACATCCTCAGACTATGC (REV). pGL3-promoter (Promega) vector (pGL3-Promoter Vector GenBank® Accession Number U47298) was used as a control to assess transfection efficiency. For transfection of the pGL3 vectors, 2 million RPE-1 cells (TxS, TxR.3 or TxR.4) were resuspended in nucleofection buffer (Solution I and II 4:1). Solution I (125 mM Na2HPO4, 12.5 mM KCl, pH 7.75) Solution II (55 mM MgCl2). After co-transfection of 100ng of Renilla plasmid (pRL-SV40 Vector GenBank® Accession Number AF025845) and 1 μg pGL3-basic-empty, 1 μg pGL3-basic-*ABCB1* or 1ug pGL3-promoter plasmid, cells were electroporated in an Amaxa 2D Nucleofector using program U-023. Cells were plated in 6-well plates and next day medium was changed. Luciferase reporter assay was performed 48h after nucleofection using a Dual-Luciferase Reporter assay kit (Promega). Cells were lysed directly on the plate with passive lysis buffer for 15 min at room temperature. Luciferase and Renilla activity were measured with the substrates from the kit using TECAN Infinite M200 PRO machine.

### Generation of CRISPRa cell lines

For RPE-1 CRISPRa, sgRNAs targeting human *ABCB1* P1, *ABCB1* P2, *ABCB4*, *RUNDC3B*, intronic regions and *POU3F2*, *LHX6* and *ZIC5* were individually cloned into the lentiCRISPR v2 plasmid. Specific sequences are found on **Sup Table 1**. CRISPR vectors were co-expressed with 3rd generation viral vectors in HEK293T cells using Fugene6 Transfection Reagent. After lentivirus production, the medium was harvested and transferred to the designated cell lines. Two days post infection cells were put on puromycin selection for two weeks.

### tracrRNA:crRNA design and transfections in RPE-1 iCut

Alt-R crRNA (Integrated DNA technologies) for LBR, LMNB1 and LMNA were obtained from the Human CRISPR Knockout Pooled Library (GeCKO v2)^62^. Specific sequences are found on **Sup Table 2**. tracrRNA:crRNA duplex was transfected according to the manufacturer’s protocol^63^.

### siRNA transfections

ON-TARGETplus SMARTpool set of 4 siRNAs targeting LBR, POU3F2, LHX6 or ZIC5 were from Dharmacon and were transfected using RNAiMAX (Life Technologies) according to manufacturer’s protocol^63^ at a final concentration of 20nM. All transfections were performed 48h before experiment, if not specified on the figure legend.

### Density and Colony Formation Assays

1 million cells were treated indicated dose of taxol and allowed to grow out for 15 days. Plates were fixed in 80% Methanol and stained with 0.2% Crystal Violet solution. Cell density was measured in ImageJ and normalized to control (WT) plate. For colony formation assays, the number of taxol resistant cells were counted.

### Viability assays

For viability assays, 1000 cells were plated in a 96-well plate and treated for 7 days with indicated drug concentrations. Subsequently, plates were fixed in 80% Methanol and stained with 0.2% Crystal Violet solution.

### RNA isolation and qRT-PCR analysis

RNA isolation was performed by using Qiagen RNeasy kit and quantified using NanoDrop (Thermo Fisher Scientific). cDNA was synthesized using Bioscript reverse transcriptase (Bioline), Random Primers (Thermo Fisher), and 1000 ng of total RNA according to the manufacturer’s protocol. Primers were designed with a melting temperature close to 60 degrees to generate 90–120-bp amplicons, mostly spanning introns. cDNA was amplified for 40 cycles on a cycler (model CFX96; Bio-Rad Laboratories) using SYBR Green PCR Master Mix (Applied Biosystems). Target cDNA levels were analyzed by the comparative cycle (Ct) method and values were normalized against GAPDH expression levels. qRT-PCR oligo sequences are summarized in **Sup Table 3**.

### Immunofluorescence

Cells were fixed with 3.7% formaldehyde and permeabilized with 0.2% Triton-X100 for 10 minutes. After, cells were blocked in 4% bovine serum albumin (BSA) in PBS supplemented with 0.1% Tween (PBS-T) for 1h. Cells were incubated for 2h at 4°C with primary antibody in PBS-T with 3% BSA, washed three times with PBS-T, and incubated with secondary antibody and DAPI in PBS-T with 3% BSA for 1h at room temperature (RT). Images were acquired with the use of a DeltaVision Elite (Applied Precision) equipped with a 60x 1.45 numerical aperture (NA) lens (Olympus) and cooled CoolSnap CCD camera. Nuclear intensity of the different chromatin marks was evaluated in ImageJ using an in-hose developed macro that enables automatic and objective analysis. The following antibodies were used for immunofluorescence experiments: H3K27ac (Actif Motif #39133, 1:500), and H3K9me2 (ab1220, 1:500). Secondary antibodies were anti-rabbit Alexa 488 (A11008 Molecular probes, 1:600), anti-mouse Alexa 568 (A11004 Molecular probes, 1:600). DAPI was used at a final concentration of 1μg/mL.

### Western Blots

For western blot experiments, equal amounts of cells were lysed with Laemmli buffer and separated by SDS–polyacrylamide gel electrophoresis followed by transfer to a nitrocellulose membrane. Membranes were blocked in 5% milk in PBST for 1h at RT before overnight incubation with primary antibody in PBST with 3% BSA at 4°C. Membranes were washed three times with PBST followed by incubation with secondary antibody in PBST with 5% milk for 2h at RT. Antibodies were visualized using enhanced chemiluminescence (ECL) (GE Healthcare). The following antibodies were used for western blot experiments: SMC1 (Bethyl, A300-055a), α-Tubulin (Sigma t5168), DNMT1 (Sigma, D4692), H3K27me3 (Actif Motif #39156), H3k27ac (Actif Motif #39133), H3k9me2 (ab1220), LaminB1 (ab16048), LaminA (sc6215) and Lamin B Receptor (ab232731). For secondary antibodies, peroxidase-conjugated goat anti-rabbit (P448 DAKO, 1:2000), goat anti-mouse (P447 DAKO, 1:2000) and rabbit anti-goat (P449) were used.

### RNA FISH

RPE-1 cells were plated on glass coverslips and washed twice with BS before fixation in 4% PFA in PBS for 10 minutes at room temperature. After two additional washes in 1x PBS coverslips were incubated in 70% ethanol at 4°C overnight. Coverslips were incubated for pre-hybridization in wash buffer (2x saline-sodium citrate (SSC) with deionized formamide (Sigma) 10%) for 2-5 minutes at room temperature. RNA FISH probe mix wash dissolved in hybridization buffer (wash buffer supplemented with 10% dextran sulfate). 38 probes labelled with Cy5 were targeted to the intronic regions of ABCB1 (Biosearch technologies). Coverslips were incubated in hybridization solution for at least 4h at 37°C. Then coverslips were washed twice for 30 minutes with wash buffer followed by a quick rinse with 2x SSC. Finally, coverslips were washed once for 5 minutes in 1x PBS before mounting on slides using Prolong gold DAPI mounting medium (Life Technologies). Images were acquired with the use of a DeltaVision Elite (Applied Precision) equipped with a 60x 1.45 numerical aperture (NA) lens (Olympus) and cooled CoolSnap CCD camera. ABCB1 transcription start site quantification was performed manually double blind.

### ChIP-sequencing of RPE-1 hTERT cells

Chromatin immunoprecipitations (ChIP) were performed as described previously^64^ with minor adjustments. For ChIP of histone marks, approximately 7.0.10^6^ million cells, 50 μL of Protein A magnetic beads (Invitrogen) and 5μg of antibody were used. Antibodies were H3K27ac (Actif Motif #39133), H3K9me3 (ab8898), H2AZ (ab4174), 5-methylcytosine (ab10805). For ChIP-seq, samples were processed for library preparation (Part# 0801-0303, KAPA Biosystems kit), sequenced using an Illumina Hiseq2500 genome analyzer (65bp reads, single end) and aligned to the Human Reference Genome (hg19) using Burrows-Wheeler Aligner (bwa) version 0.5.9. Mapped reads were filtered based on mapping quality of 20 using samtools version 0.1.19. For ChIP-qPCR analysis, DNA was amplified for 40 cycles on a cycler (model CFX96; Bio-Rad Laboratories) using SYBR Green PCR Master Mix (Applied Biosystems). Target DNA levels were analyzed by the comparative cycle (Ct) method and values were normalized against input DNA and positive control region (specific for each chromatin mark). ChIP-qPCR oligo sequences are summarized in **Sup Table 3**.

### RNA-sequencing

Total RNA from cultured cells was extracted using RLT (Quiagen). Strand-specific libraries were generated using the TruSeq PolyA Stranded mRNA sample preparation kit (Illumina). In brief, polyadenylated RNA was purified using oligo-dT beads. Following purification, the RNA was fragmented, random-primed and reserve transcribed using SuperScript II Reverse Transcriptase (Invitrogen). The generated cDNA was 3′ end-adenylated and ligated to Illumina Paired-end sequencing adapters and amplified by PCR using HiSeq SR Cluster Kit v4 cBot (Illumina). Libraries were analyzed on a 2100 Bioanalyzer (Agilent) and subsequently sequenced on a HiSeq2000 (Illumina). We performed RNAseq alignment using TopHat 2.1.1. Differentially expressed genes were called with DEseq2, with an adjusted p-value threshold of 0.05.

### TLA analysis

TLA was performed as previously described with minor modifications^39^. TLA libraries were sequenced on a MiSeq and were analyzed with a custom TLA mapping pipeline. TLA ligation data were mapped to hg19. Normalization and downstream analysis were done using peakC16.

### DamID-seq

DamID-seq was performed as described^65^ with minor modifications. Dam fused to human LMNB1 protein (Dam-LMNB1) or unfused Dam were expressed in cells by lentiviral transduction^66^. Three days after infection, cells were collected for genomic DNA (gDNA) isolation. gDNA was pre-treated with SAP (10 U, New England Biolabs #M0371S) in CutSmart buffer in a total volume of 10 μl at 37°C for 1h, followed by heat-inactivation at 65°C for 20 min to suppress signal from apoptotic fragments. This gDNA was then digested with DpnI (10 U, New England Biolabs #R0176L) in CutSmart buffer in a total volume of 10 μl at 37°C for 8h followed by heat inactivation at 80 °C for 20 min. Fragments were ligated to 12.5 pmol DamID adapters using T4 ligase (2.5 U, New England Biolabs ##) in T4 ligase buffer in a total volume of 20 μl incubated at 16°C for 16h. The reaction was heat-inactivated for 10 minutes at 65°C. Products were then digested with DpnII to destroy partially methylated fragments. DpnII buffer and DpnII (10 U, New England Biolabs #R0543L) were added in a total volume of 50 μl and incubated at 37 °C for 1 h. Next, 8 μl of DpnII-digested products was amplified by PCR with MyTaq Red Mix (Bioline #BIO-25044) and 1.25 μM primers Adr-PCR-Rand1 in a total volume of 40 μl. PCR settings were 8 min at 72 °C (1×) followed by 20 s at 94 °C, 30 s at 58 °C, 20 s at 72 °C (24× for Dam, 28x for Dam-LMNB1 samples) and 2 minutes at 72°C (1×). Remaining steps were performed as previously described. Samples were sequenced on an Illumina HiSeq2500.

### Motif analysis

Genomic coordinates of all the genes were obtained from GRCh37 (Ensembl version 75) using biomaRt package^67^ and transcription starting sites of the genes were extend 1 kb to both up- and down-stream to identify the promoter regions. The motifs presenting in the promoters were identified using GimmeMotifs^68^ against the non-redundant Cis-bp database (version 3.0). To identify the overrepresented motifs, we used a similar method as described in our previous publication (https://www.nature.com/articles/s41588-020-00744-4, will be online next Monday). Briefly, we calculated for every motif the frequency in the promoters of the upregulated genes and all the expressed genes. We computed relative motif frequency by dividing the individual motif frequency by to total number of identified motifs. We calculated the log2-enrichment score by calculating the ratio of relative motif frequency between the promoters of up-regulated genes and all the expressed genes. The p-value was calculated using the Fisher exact test on the following 2×2 table: for every motif M, we determine the number of the promoters belonging to the upregulated genes with or without M and for the promoters of the expressed genes with or without M.

### Processing of RPE-1 DamID data

DamID-seq was performed as described in ^44^

---

**Supplementary Table 1.**
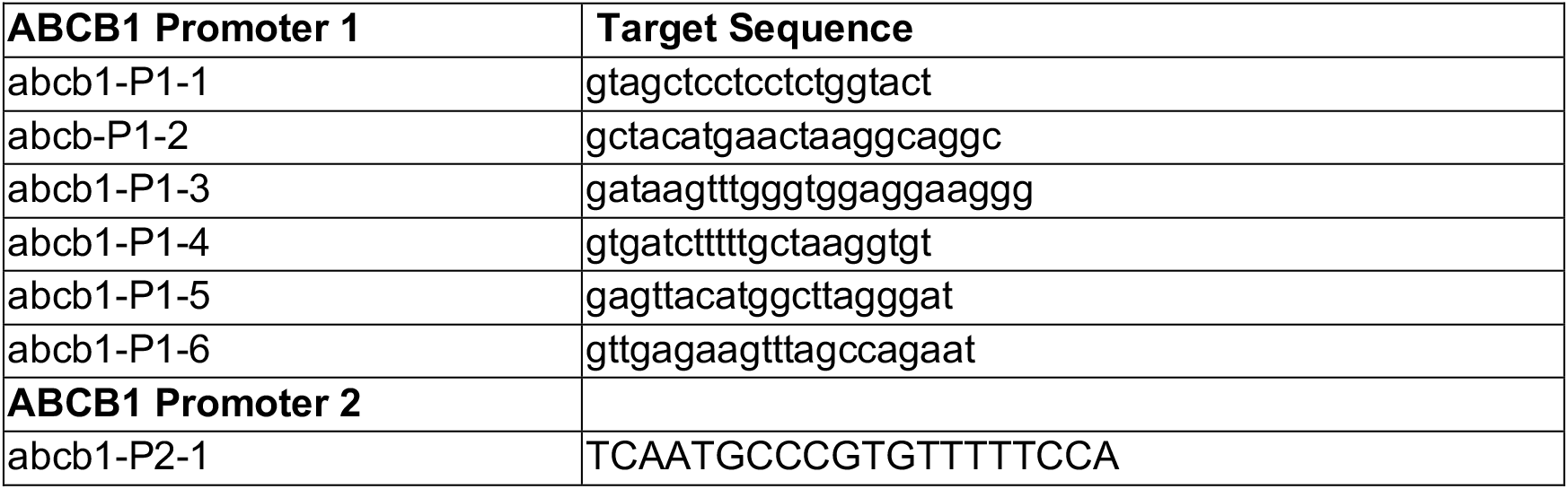
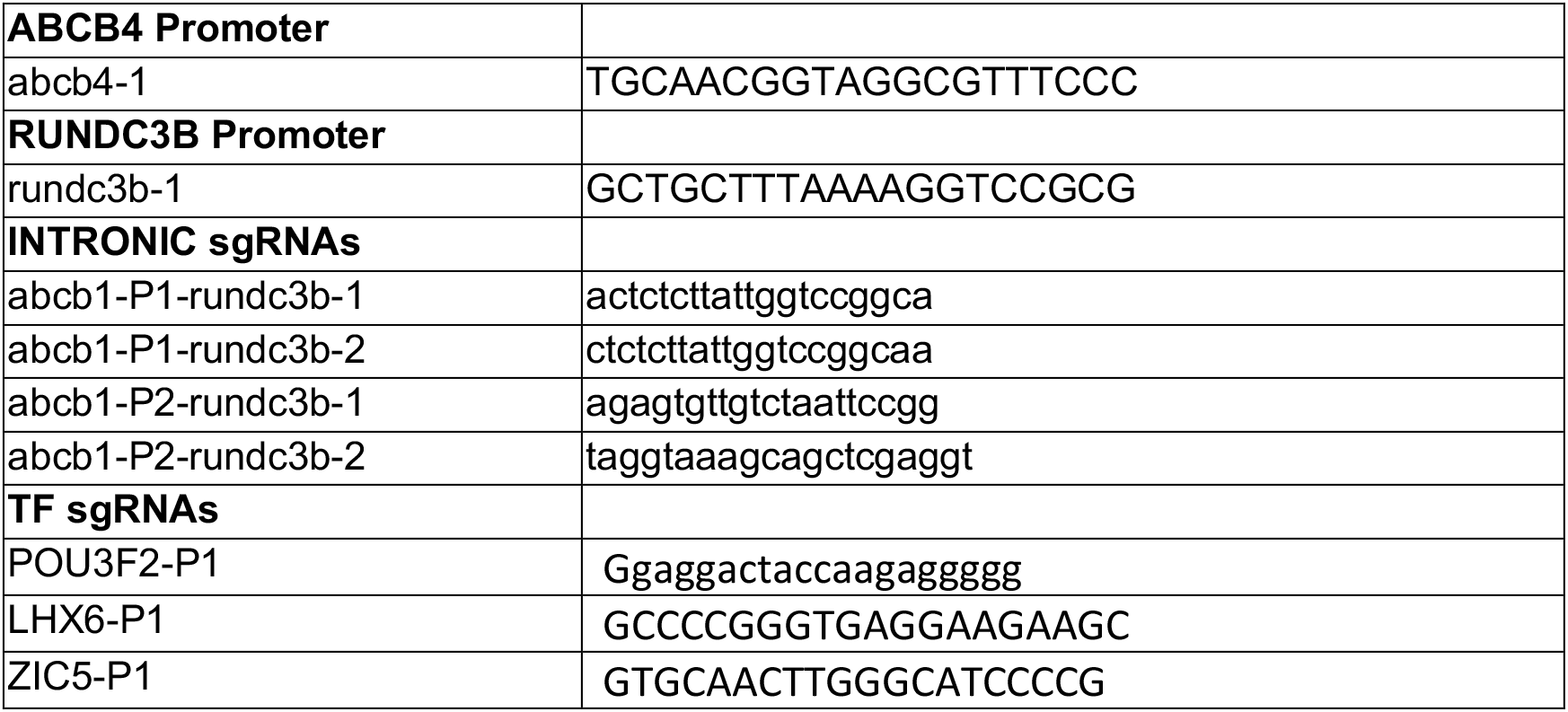
sgRNAs for RPE-1 CRISPRa

**Supplementary Table 2.**
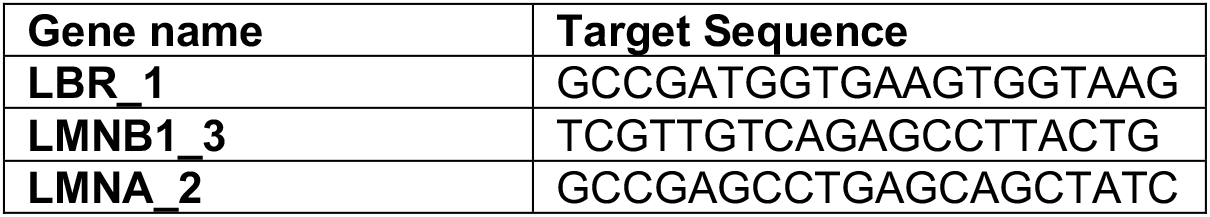
crRNA for RPE-1 icut KO

**Supplementary Table 3.**
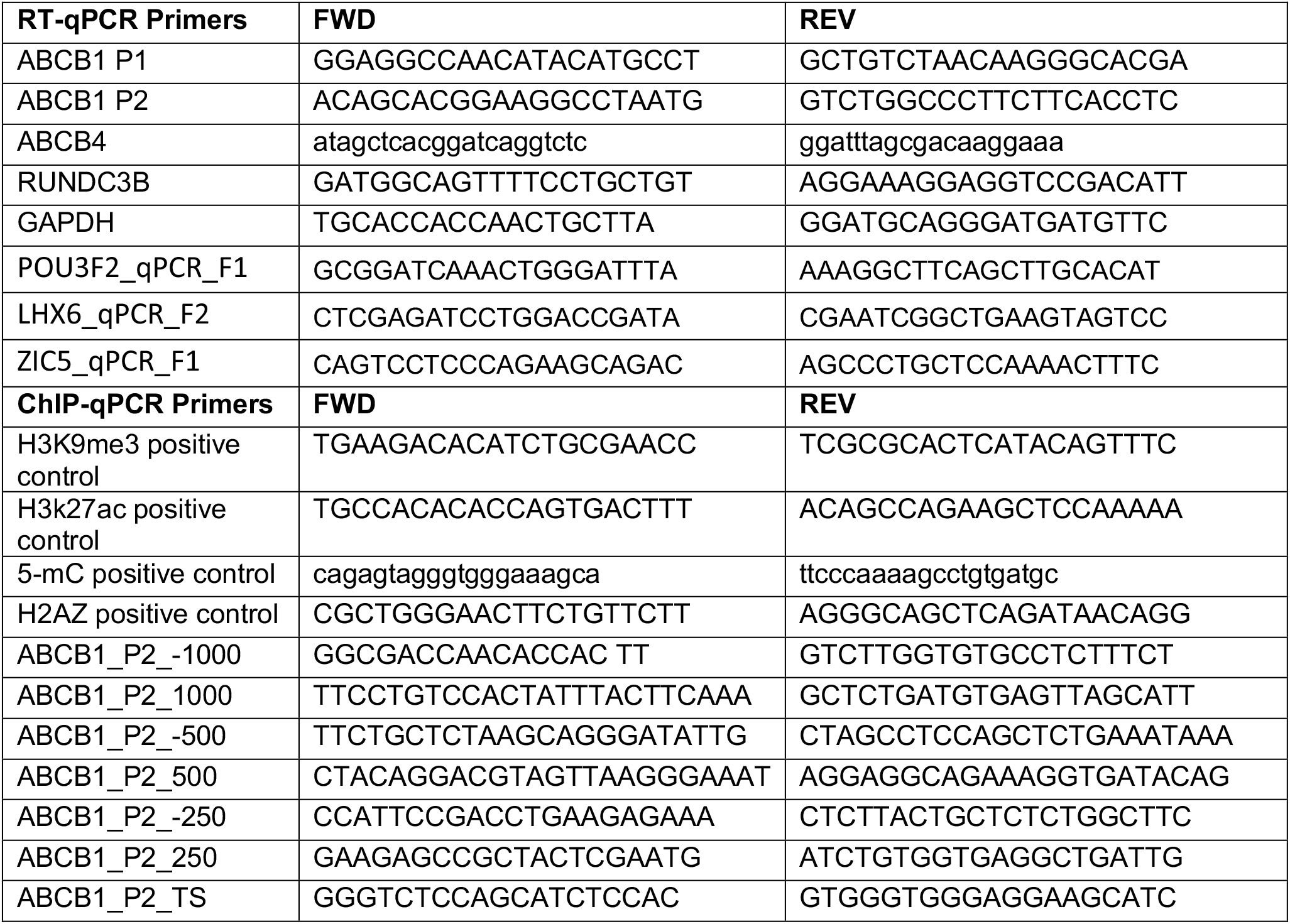
RT-qPCR and ChiP-qPCR primers

**Supplementary Table 4.**
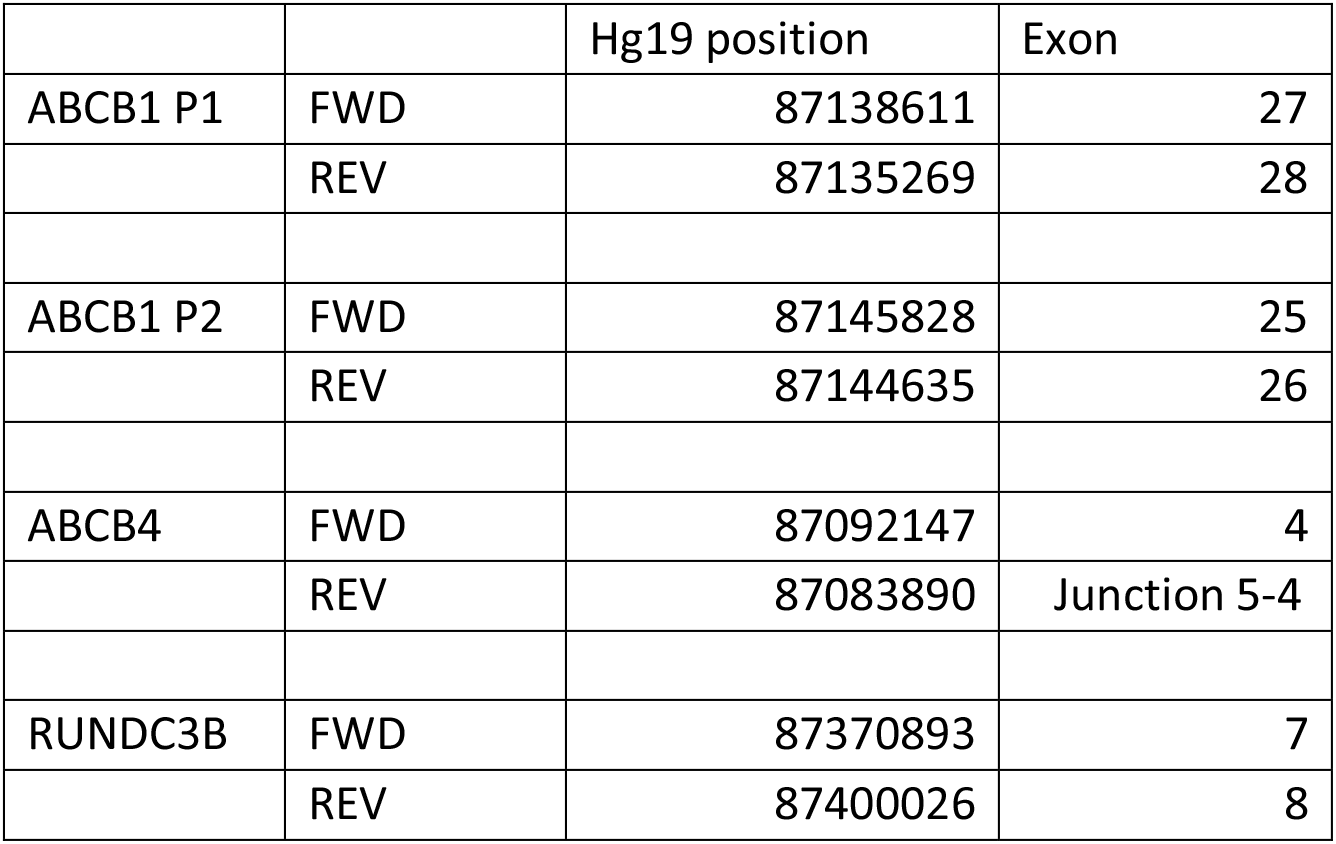
RT-qPCR primers position

## Supplementary figure legends

**Supplementary Figure 1.**
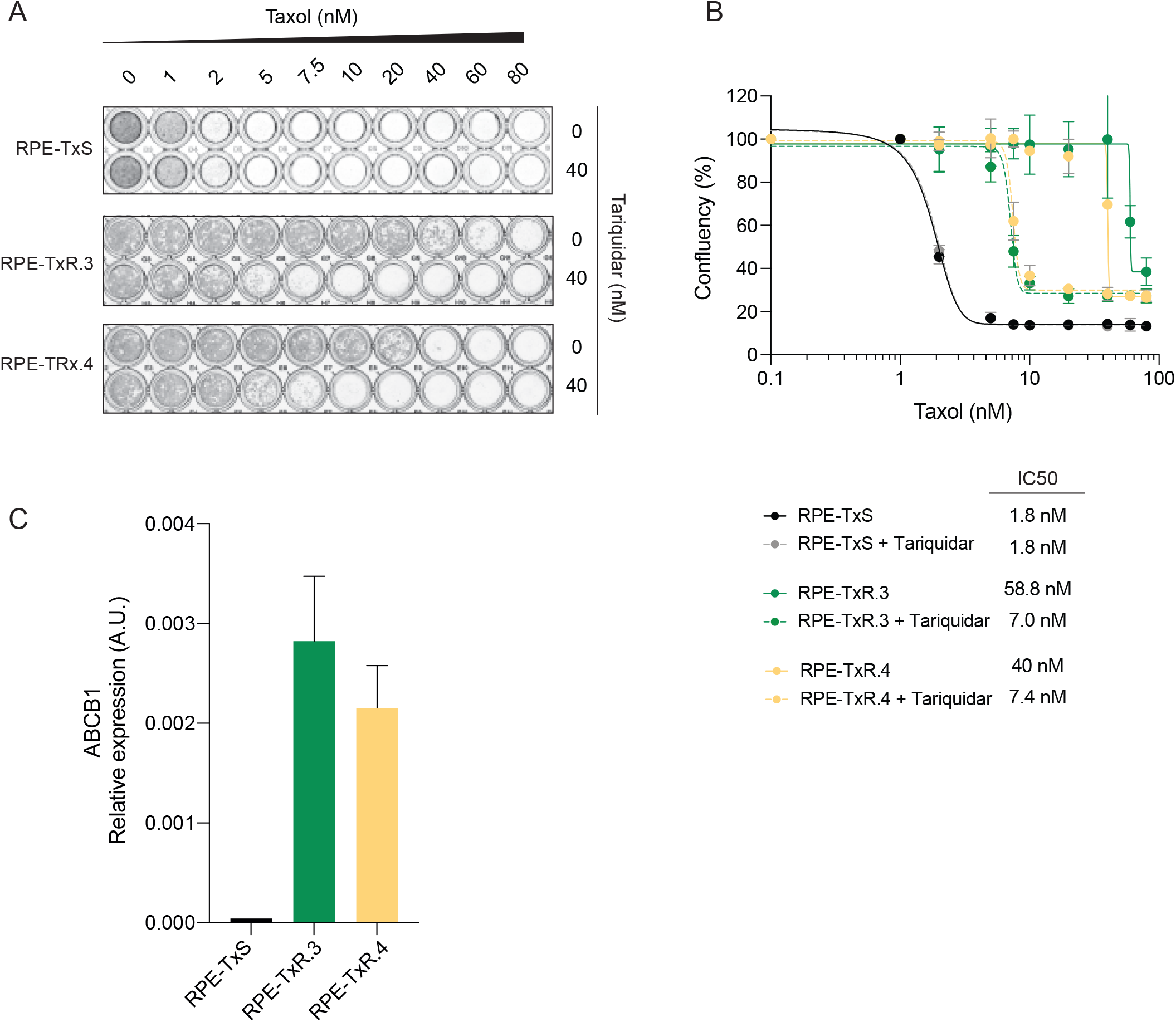
Transcriptional activation of *ABCB1* drives taxol resistance in independently generated RPE-TxR. **A)** Crystal violet staining of viability assay on taxol-naïve RPE-TxS and two independently generated taxol resistant cell lines (RPE-TxR3 and RPE-TxR4). **B)** Relative survival plots of the RPE-TxS and RPE-TxR3 and TxR4 cell lines. Error bars show the average +/− s.d. of two independent experiments and the calculated IC50. The curve was drawn from the log(inhibitor) vs response equation *Y=Bottom + (Top-Bottom)/(1+10^(X-LogIC50))*. **C)***ABCB1* mRNA levels determined by qRT-PCR and normalized to *GAPDH* expression in RPE-TxS, RPE-TxR3 and RPE-TxR4, n=2. Error bars show the SD.

**Supplementary Figure 2.**
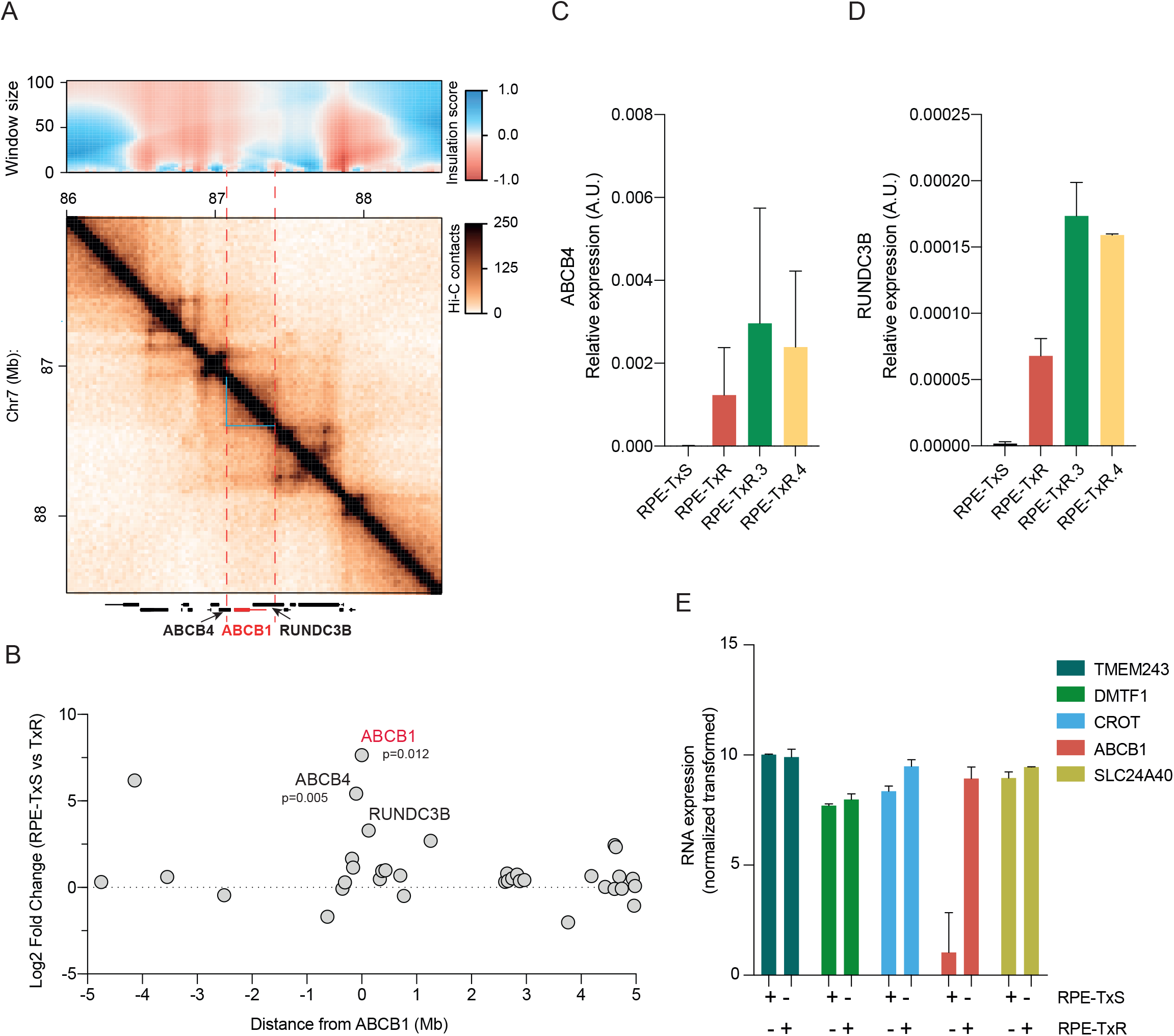
RPE-TxR undergo changes in gene expression. **A)** Hi-C contact matrix of RPE-1 WT generated by Aiden Lab. For TAD calling, we calculated the insulation score for each bin at 25kb resolution using the software GENOVA^69^. Blue lines show the TAD called where *ABCB1* is located together with *ABCB4* and *RUNDC3B*. **B)** Log2 Fold change of RNA expression levels of genes across 5Mb +/− *ABCB1* comparing RPE-TxS to RPE-TxR, n=2. Every dot indicates a gene. **C)***ABCB4* and **D)***RUNDC3B* mRNA levels determined by qRT-PCR and normalized to *GAPDH* expression in RPE-TxS, RPE-TxR, RPE-TxR3 and RPE-TxR4, n=2. Error bars show the SD. **E)** Normalized RNA expression of *ABCB1* and its neighbor transcribed genes in RPE-TxS and RPE-TxR cells.

**Supplementary Figure 3.**
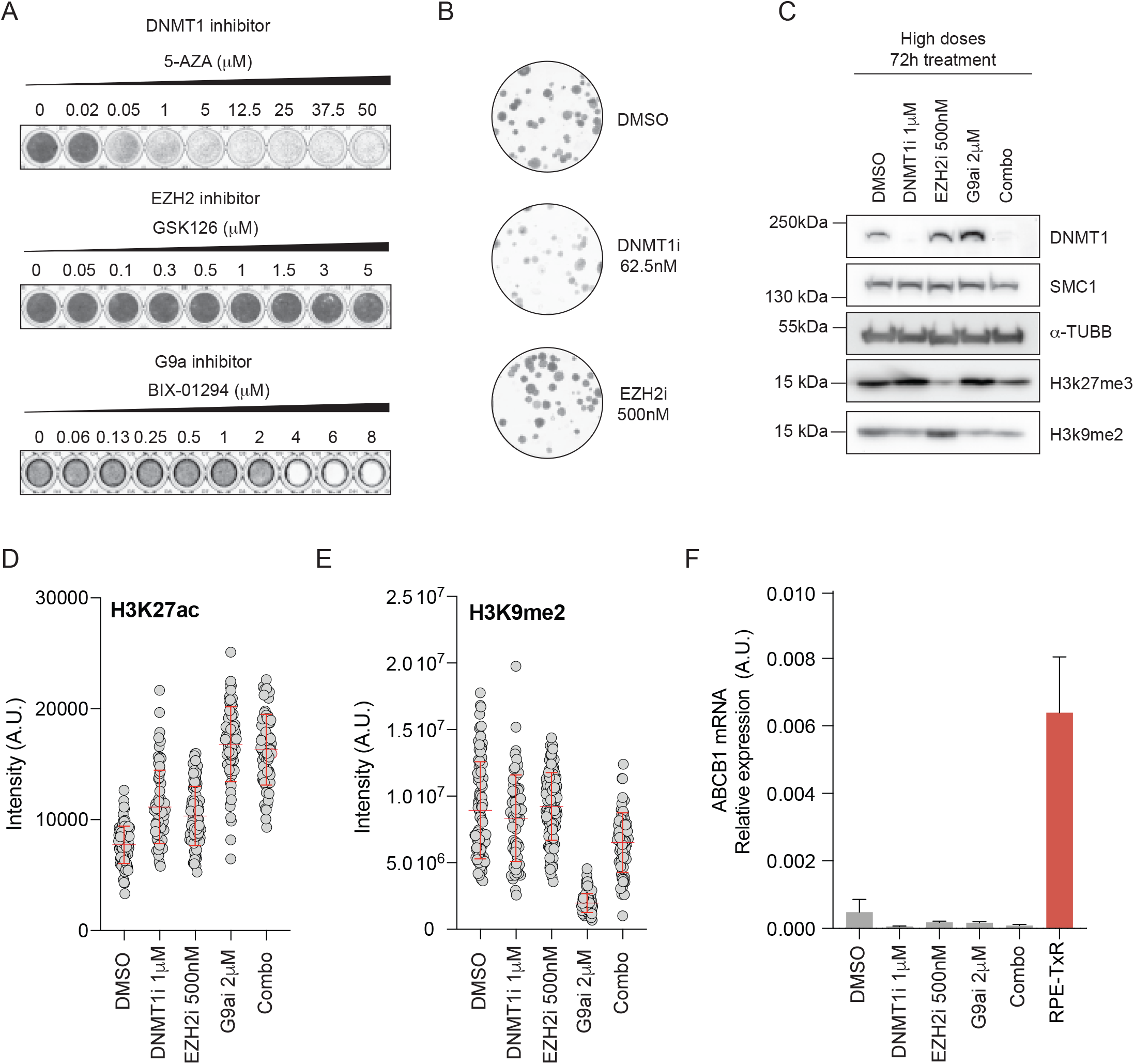
5-AZA and GSK126 inhibitors validations. **A)** Crystal violet staining of viability assay on RPE-TxS with increasing concentration of the epigenetic drugs 5-Aza-2’-deoxycytidine (5-AZA, DNMT1 inhibitor), GSK126 (EZH2 inhibitor) and BIX-01294 (G9a inhibitor). **B)** Crystal violet staining of colony formation assay under the indicated drug doses without taxol. 100 cells were plated per condition and let grown for 15 days in parallel to Fig 3C. **C)** Western Blot showing the levels of the chromatin proteins and controls (SMC1 an α-TUBB) upon treatment with single drugs or the combination (Combo) for 72h. **D)** Immunofluorescence quantification of nuclear H3K27ac and **E)** H3K9me2 levels after 72h drug addition by ImageJ in-house foci macro, n=1, 60 cells per condition. Error bars show the SD. **F)***ABCB1* mRNA levels determined by qRT-PCR and normalized to *GAPDH* expression levels upon high drug addition and RPE-TxR as a control for *ABCB1* expression, n=3 technical replicates. Error bars show the SD.

**Supplementary Figure 4.**
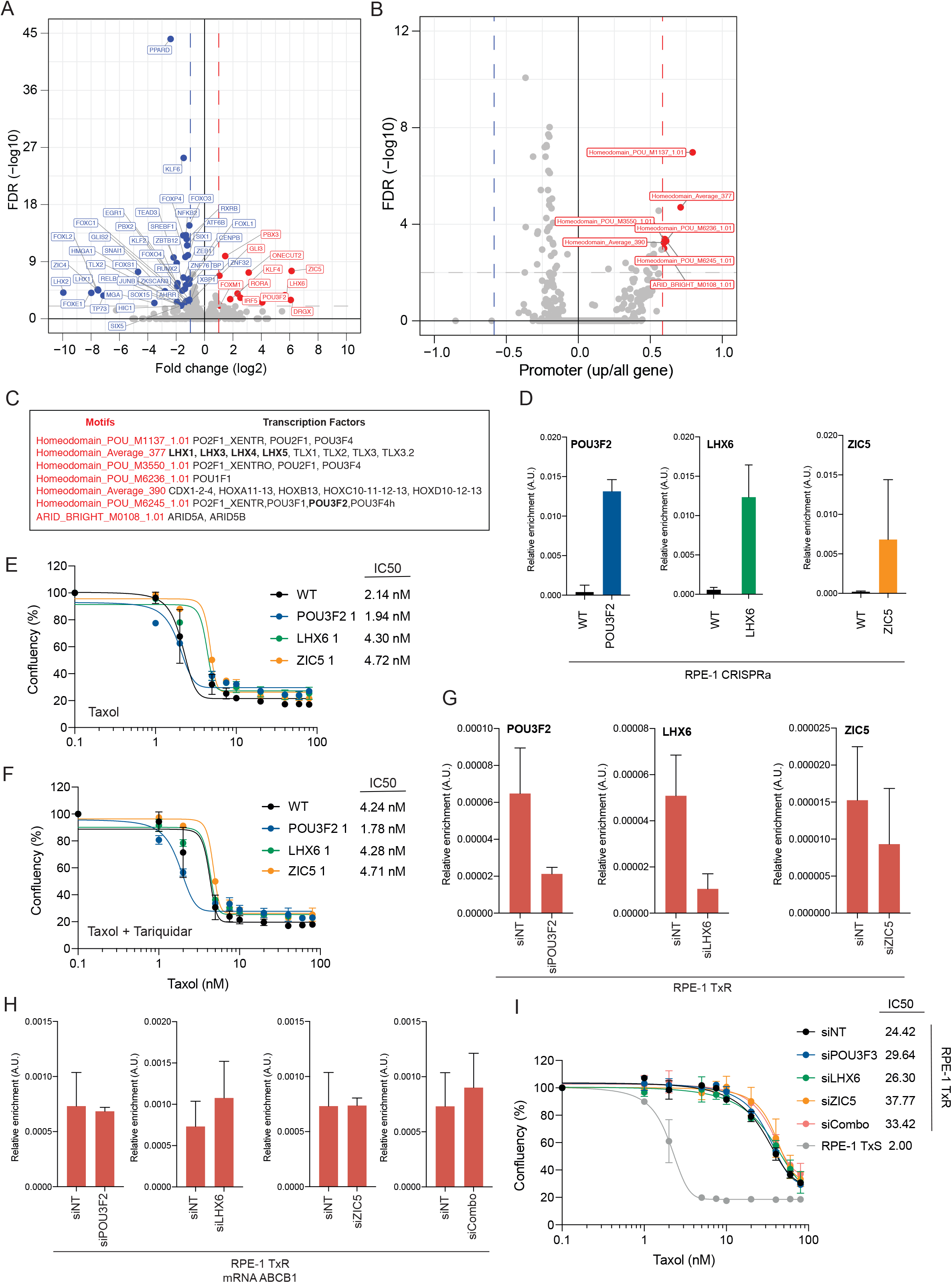
*POU3F2*, *LHX6* or *ZIC5* are not responsible for initiation of *ABCB1* gene expression in RPE-TxR cells. **A)** RNA-seq analysis identified the differentially regulated transcription factors genes in the taxol resistant RPE-TxR cells (n=2) compared to RPE-TxS cells (n=3). **B)** Motif analysis revealed the potential promoter activators in the RPE-TxR cell line. **C)** Table showing the corresponding TF binding the significant motifs found on B. **D)** mRNA levels determined by qRT-PCR and normalized to *GAPDH* expression in RPE-1 CRISPRa targeted with sgRNA for *POU3F2*, *LHX6* or *ZIC5*, n=2. Error bars show the SD. **E and F)** Relative survival plots of the same TFs CRISPRa cell lines. Error bars show the average +/− s.d. of two independent experiments and the calculated IC50. The curve was drawn from the log(inhibitor) vs response equation *Y=Bottom + (Top-Bottom)/(1+10^(X-LogIC50))*. **G)** mRNA levels of the TF candidates or **H)***ABCB1* determined by qRT-PCR and normalized to *GAPDH* expression in RPE-TxR transfected with siRNA NT, siPOU3F2 or siZIC5. Error bars show the SD, n=2. **I)** Relative survival plots of the respective siRNA transfections. Error bars show the average +/− s.d. of two independent experiments and the calculated IC50.

**Supplementary Figure 5.**
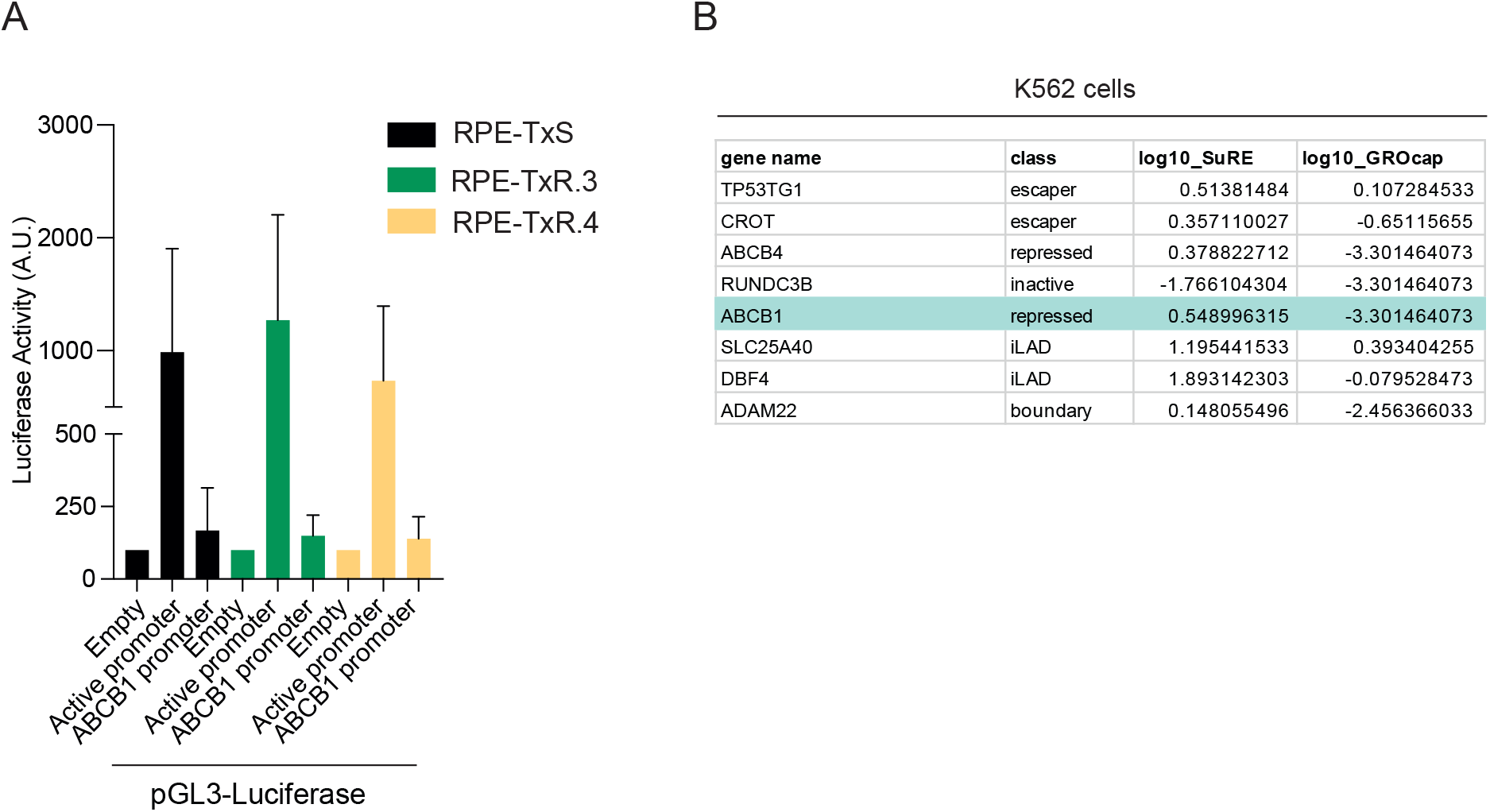
*ABCB1* upregulation in RPE-TxR is not caused by direct activation of the promoter by trans-acting factors. **A)** Relative luciferase activity calculated by dividing the luciferase activity to that of Renilla luciferase. Data shown represent average +/− s.d, n = 3. **B)** Promoter classification of the genes neighboring ABCB1 based on GROcap and SuRE in K562 cells.

**Supplementary Figure 6.**
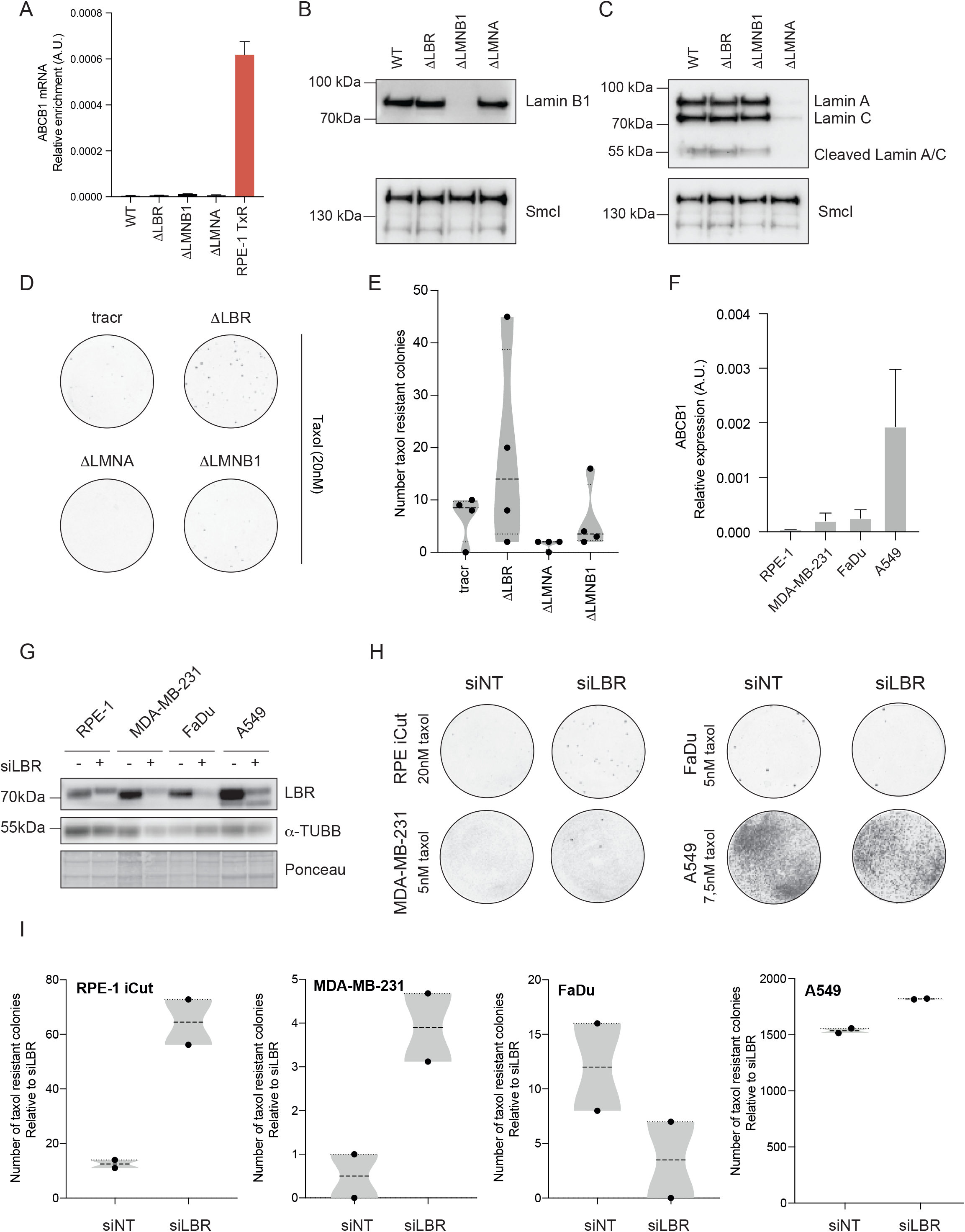
*LBR*, *LMNB1* and *LMNA* knockout and knockdown validations. **A)** ABCB1 mRNA levels determined by qRT-PCR and normalized to GAPDH expression in RPE-iCut in WT cells or 7 days after transfection of *LBR*, LMNA or LMNB1 crRNAs. n=2. Error bars show the SD. **B)** Western Blot showing Lamin B1 and control (SMC1) protein levels upon the different KO in RPE-1 iCut cells. **C)** Western Blot showing LaminA/C and control (SMC1) protein levels upon the different KO in RPE-1 iCut cells. **D)** Crystal violet staining of colony formation assay under 20nM of taxol in RPE-1 iCut WT, transfected only with tracrRNA or together with the specific crRNA to generate a KO. **E)** Quantification of the number of taxol resistant colonies under 20nM of taxol in the different KO, n=2. Error bars show the SD. Black dots show an independent biological replicate. **F)** ABCB1 mRNA levels determined by qRT-PCR and normalized to GAPDH in RPE-1 and the different cancer cell lines, n=3. Error bars show the SD of technical replicates. **G)** Western Blot showing *LBR* and control (α-TUBB) protein levels in the different cancer cell lines upon *LBR* siRNA depletion. **H)** Crystal violet staining of colony formation assay in RPE-1 iCut and cancer cell lines under indicated concentration of taxol. Cells were treated for 72hrs prior to colony formation plating with siNT or *siLBR*. **I)** Quantification of the number of taxol resistant colonies in G. Black dots show an independent biological replicate.

**Supplementary Figure 7.**
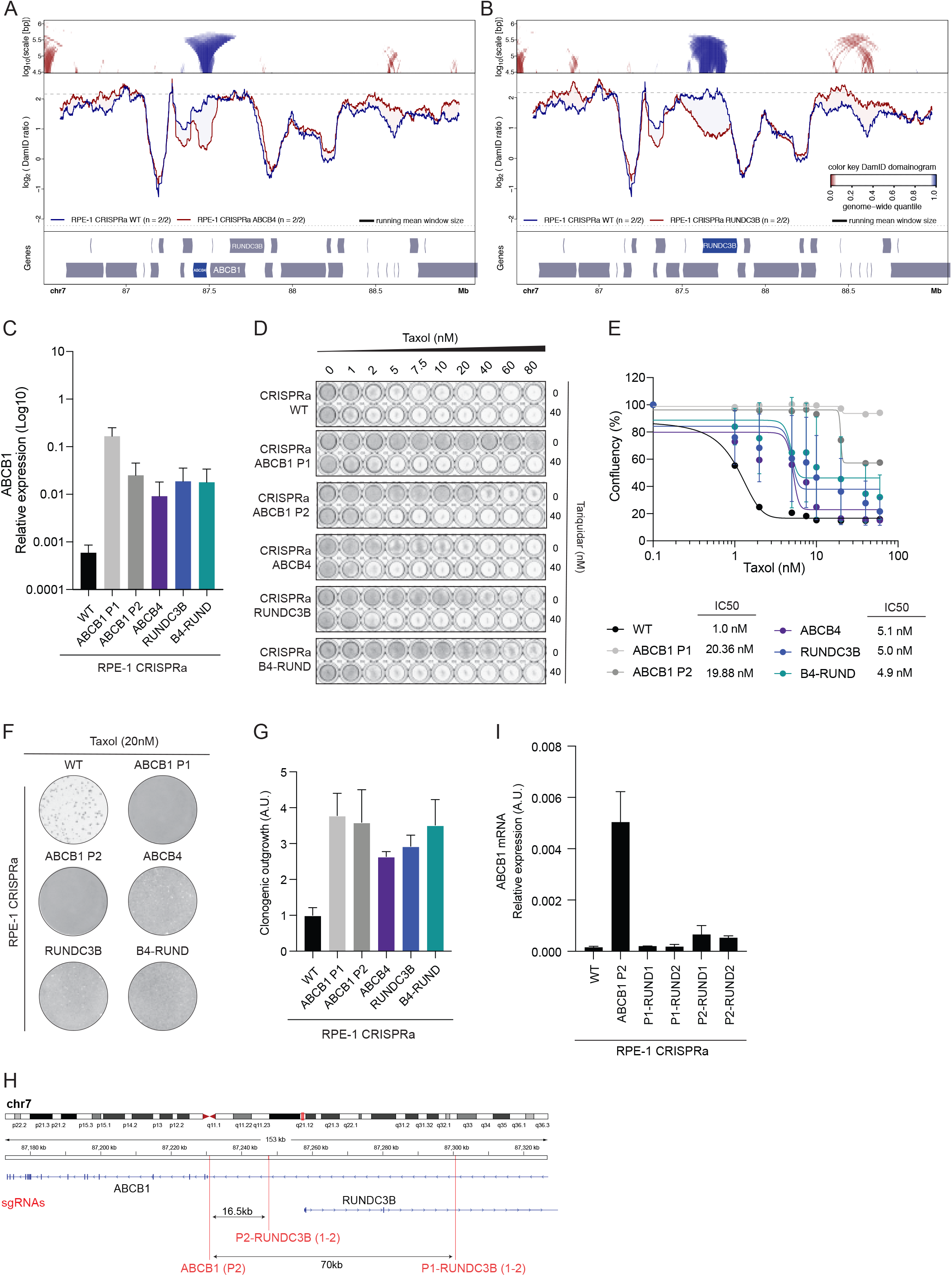
Validation of CRISPRa cell lines. **A)** Local NL detachment caused by ABCB4 gene activation or **B)** RUNDC3B by CRISPRa in RPE-1 cells. **C)** ABCB1 mRNA levels determined by qRT-PCR and normalized to GAPDH upon CRISPRa activation of individual genes or combination of ABCB4 and RUNDC3B (B4-RUND), n=2. Error bars show the SD. **D)** Crystal violet staining of viability assay on CRISPRa cell lines upon activation of ABCB1 (P1 and P2), ABCB4, RUNDC3B or the combination. **E)** Relative survival plots of CRISPRa cell lines targeting ABCB1 (P1 and P2), ABCB4, RUNDC3B or the combination of the last two. Error bars show the average +/− s.d. of two independent experiments and the calculated IC50. The curve was drawn from the log(inhibitor) vs response equation *Y=Bottom + (Top-Bottom)/(1+10^(X-LogIC50))*. **F)** Crystal violet staining of density assays of CRISPRa cells targeting the different genes upon 20nM of taxol. WT, ABCB1 P2 and B4-RUND are duplicated from Figure 5E. **G)** ImageJ quantification of density assays of CRISPRa cells targeting the different genes upon 20nM of taxol, n=2. Error bars show the SD. **H)** Schematic representation of the Chr7q21.12 region indicating the locations where the sgRNAs were targeting intronic regions for the ABCB1 gene. **I)** ABCB1 mRNA levels determined by qRT-PCR and normalized to GAPDH in CRISPRa cells upon sgRNA targeting of the different intronic regions, n=2. Error bars show the SD.

